# The structural basis of Indisulam-mediated recruitment of RBM39 to the DCAF15-DDB1-DDA1 E3 ligase complex

**DOI:** 10.1101/737510

**Authors:** Dirksen E. Bussiere, Lili Xie, Honnappa Srinivas, Wei Shu, Ashley Burke, Celine Be, Junping Zhao, Adarsh Godbole, Dan King, Rajeshri G. Karki, Viktor Hornak, Fangmin Xu, Jennifer Cobb, Nathalie Carte, Andreas O. Frank, Alexandra Frommlet, Patrick Graff, Mark Knapp, Aleem Fazal, Barun Okram, Songchun Jiang, Pierre-Yves Michellys, Rohan Beckwith, Hans Voshol, Christian Wiesmann, Jonathan Solomon, Joshiawa Paulk

**Affiliations:** Novartis Institutes for Biomedical Research, Emeryville, CA, USA; Novartis Institutes for Biomedical Research, Basel, Switzerland; Novartis Institutes for Biomedical Research, Cambridge, MA, USA; Genomics Institute of the Novartis Research Foundation, San Diego, CA, USA

## Abstract

The anti-cancer agent Indisulam inhibits cell proliferation by causing degradation of RBM39, an essential mRNA splicing factor. Indisulam promotes an interaction between RBM39 and the DCAF15 E3 ligase substrate receptor leading to RBM39 ubiquitination and proteasome-mediated degradation. To delineate the precise mechanism by which Indisulam mediates DCAF15-RBM39 interaction, we solved the DCAF15-DDB1-DDA1-Indisulam-RBM39(RRM2) complex structure to 2.3 Å. DCAF15 has a novel topology which embraces the RBM39(RRM2) domain largely via nonpolar interactions, and Indisulam binds between DCAF15 and RBM39(RRM2) and coordinates additional interactions between the two proteins. Studies with RBM39 point mutants and Indisulam analogs validated the structural model and defined the RBM39 alpha-helical degron motif. The degron is found only in RBM23 and RBM39 and only these proteins were detectably downregulated in Indisulam-treated HCT116 cells. This work further explains how Indisulam induces RBM39 degradation and defines the challenge of harnessing DCAF15 to degrade novel targets.

## Introduction

Targeted protein degradation (TPD) is an emerging area of small molecule drug discovery^12^. In TPD, small molecules do not directly modulate the activity of their target proteins upon binding, but instead bring about the interaction of targets with E3 ligases of the Ubiquitin-Proteasome System (UPS). This compound-induced proximity of the target and E3 ligase leads to removal of the target protein from the cell by proteolytic degradation.

The Ubiquitin-Proteasome System exists in every cell and functions to regulate most protein half-life^3^. Conjugation of four or more copies of ubiquitin, a small 76-amino acid protein, allows protein recognition by the 26S proteasome^4^. Upon binding to the proteasome lid, poly-ubiquitinated proteins are pulled into the proteasome tube and cleaved by interior proteolytic active sites into peptide fragments^56^. Ubiquitination is tightly regulated by a three enzyme cascade^7^. Ubiquitin is activated by the E1 enzyme and is transferred to one of the E2 enzymes. The E3 ligases determine which proteins are mono- or poly-ubiquitinated by catalyzing the transfer of ubiquitin from an E2 enzyme to a lysine residue on the target protein or ubiquitin. There are over 600 E3 ligases encoded in the human genome allowing for the recognition and regulation of large number of diverse substrates, although the structural features recognized (known as the ‘degron’) are unknown for most of these ligases.^8^

In TPD, small molecules are used to hijack the E3 ligases of the UPS by a variety of mechanisms. The selective estrogen receptor degraders (SERDs) bind and destabilize the estrogen receptor (ER), increasing its surface hydrophobicity^9^. SERD-bound ER is recognized as unfolded by the protein quality control pathway and is degraded by the UPS^10^. Bifunctional degraders are modular molecules that have an E3-binding moiety, a linker, and a target-binding moiety^11^. Bifunctional degraders literally tether target proteins to E3 ligases to facilitate ubiquitination and degradation. Auxin, a small molecule phytohormone, binds to an E3 ligase forming a new ligase binding surface with increased affinity for the target protein^12^. Because auxin was described as a “molecular glue”^13^, this type of TPD molecule, is known as a molecular glue degrader. The IMiD drugs were recently discovered to be molecular glue degraders. They bind the CRBN E3 ligase and create a new binding surface that recruits beta-hairpin containing proteins^14^. Another class of TPD molecule is described by the plant hormone Gibberellin (GA). GA binds to its receptor and induces a conformational change that allows receptor binding to its target protein. The receptor-GA-target protein complex is recognized by the E3 ligase leading to target protein degradation^12^.

Indisulam (Fig. 1a), an anti-cancer agent, was recently found to be a TPD molecule. Originally discovered by screening sulfonamides for cancer cell growth inhibition,^15^ Indisulam stood out by causing G1/S cell cycle arrest and demonstrating efficacy in multiple tumor xenograft models^16^. Two seminal papers revealed that Indisulam inhibits cell growth by degrading the essential splicing factor RBM39^17, 18^. Indisulam mediates an interaction between RBM39 and the E3 ligase DCAF15 leading to RBM39 poly-ubiquitination and proteasomal degradation. It was unclear whether Indisulam acts allosterically by binding DCAF15 or RBM39 to bring about a conformational change that enhances DCAF15-RBM39 interaction, whether Indisulam stabilizes a weak DCAF15-RBM39 interaction, or whether Indisulam acts as a molecular glue to enhance RBM39 binding to DCAF15 (Fig. 1a).

**Figure 1:**
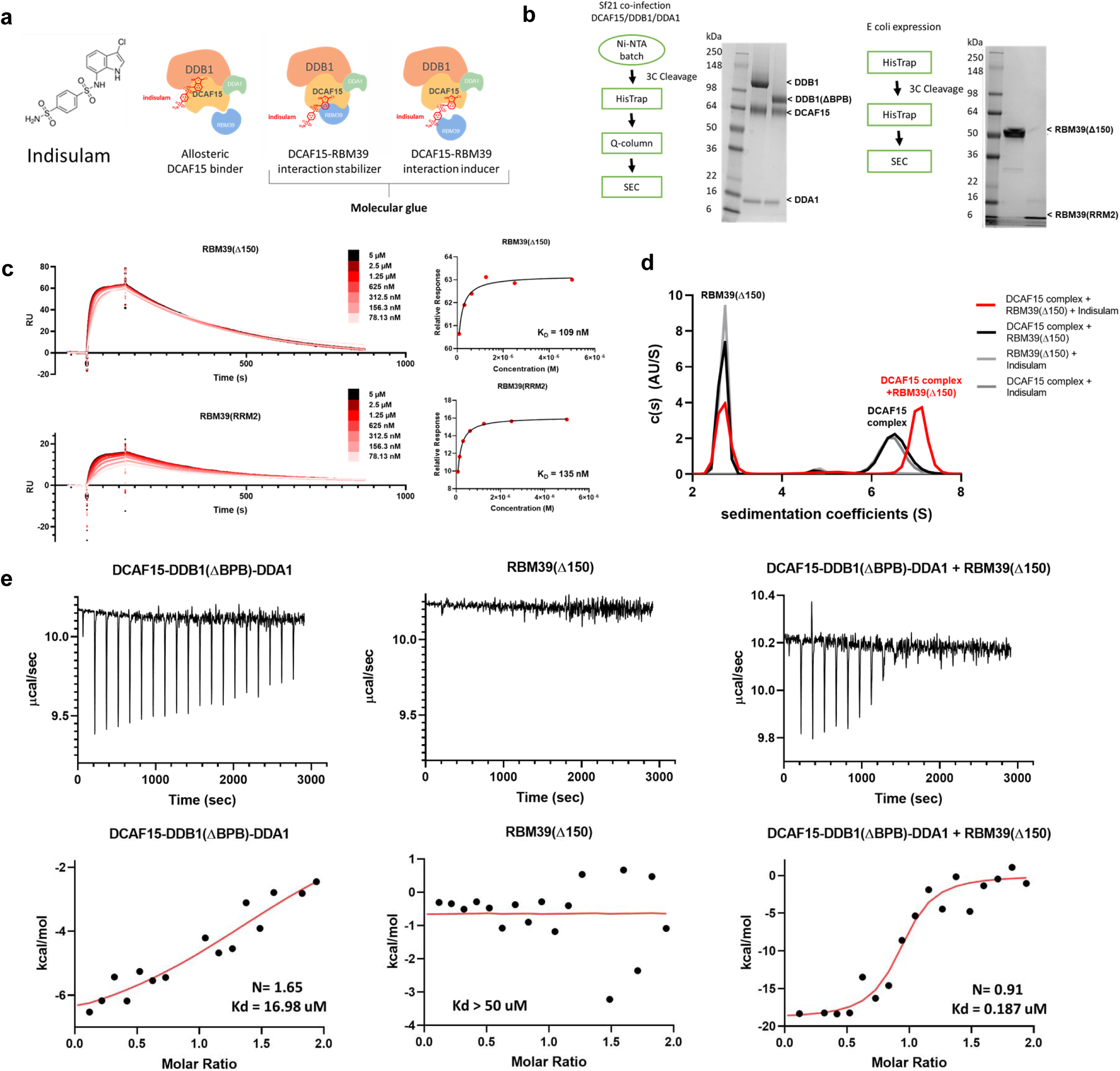
Purification and functional validation of RBM39 and DCAF15-DDB1-DDA1 complexes. **a**, Structure of Indisulam and representations of potential mechanisms of action for Indisulam-mediated recruitment of RBM39 to DCAF15. **b,** Expression systems, purification workflows, and Coomassie-stained gels for DCAF15-DDB1-DDA1, DCAF15-DDB1(ΔBPB)-DDA1 (left), RBM39(Δ150), and RBM39(RRM2) (right) proteins. **c,** Surface plasmon resonance (SPR) data characterizing Indisulam-mediated interaction between purified DCAF15-DDB1-DDA1 and RBM39 proteins. Biotinylated-DCAF15-DDB1-DDA1 was captured on the surface of SA chip and response was measured following injection of varied concentrations of RBM39(Δ150) (top) or RBM39(RRM2) (bottom) in the presence of 20 µM Indisulam. This experiment was repeated two independent times with representative data shown. **d,** Analytical ultracentrifugation (AUC) analysis of the interaction between 2.5 µM DCAF15-DDB1(ΔBPB)-DDA1 (DCAF15 complex) and 10 µM His-ZZ-RBM39(Δ150) in presence (red line) and absence (black line) of Indisulam. This experiment was repeated two independent times with representative data shown. **e,** Isothermal calorimetry measurements on 50 µM DCAF15-DDB1(ΔBPB)-DDA1 (left), 50 µM RBM39(Δ150) (middle), and a mixture of 10 µM both proteins (right) upon injections of 500 µM or 100 uM Indisulam. Corresponding fits reveal K_d_ measurements for Indisulam to be 17 µM for DCAF15-DDB1(ΔBPB)-DDA1 alone, >50 µM for RBM39(Δ150) alone, and 187 nM for DCAF15-DDB1(ΔBPB)-DDA1 and RBM39(Δ150) mixture. Representative data shown from an experiment performed two independent times and once for DCAF15-DDB1(ΔBPB)-DDA1-Indisulam experiment.

In this work, we set out to understand the precise molecular mechanism by which Indisulam brings about the interaction between RBM39 and the DCAF15 E3 ligase substrate receptor. DCAF15-DDB1-DDA1-Indisulam-RBM39 complex structures were determined by both X-ray crystallography and cryogenic electron microscopy (cryo-EM) to 2.3 and 3.5 Å, respectively. The structures reveal that Indisulam is a molecular glue degrader that binds to DCAF15 creating a novel ligase surface that enhances RBM39 binding. This detailed understanding of Indisulam’s mechanism of action is the first step towards determining whether the DCAF15 E3 ligase can be reprogrammed by other small molecules to degrade novel targets beyond RBM39.

## Purification and Characterization of the DCAF15-DDB1-DDA1-Indisulam-RBM39 complex

The DCAF15-DDB1-DDA1 complex was expressed and purified from SF21 insect cells (Fig. 1b). RBM39(Δ150) and the second RRM domain of RBM39, RBM39(RRM2), a domain implicated in Indisulam resistance ^17, 18^, were expressed and purified from *E. Coli* (Fig.1b; Supplementary Fig. 1). The purified proteins were functionally validated by measuring whether Indisulam-mediated interactions could be detected. In SPR studies, biotinylated DCAF15-DDB1-DDA1 was immobilized on the sensor surface and the response to increasing concentrations of RBM39(Δ150) or RBM39(RRM2) was measured in the presence of 20 µM Indisulam (Fig. 1c). Both RBM39(Δ150) and RBM39(RRM2) bound DCAF15-DDB1-DDA1 in an Indisulam-dependent manner with similar affinities (Kds of 109 and 135 nM, respectively). This data suggests that RBM39(RRM2) is sufficient to engage the DCAF15 complex in the presence of Indisulam. DCAF15 complex interaction with RBM39 was also interrogated by analytical ultracentrifugation (AUC). An interaction between the DCAF15-DDB1(ΔBPB)-DDA1 complex and RBM39(Δ150) was detected in the presence of Indisulam, but not with a DMSO vehicle control (Fig. 1d). In the absence of Indisulam, no interaction between the DCAF15 complex and RBM39 could be detected by AUC even upon increasing RBM39(Δ150) concentrations up to 80 µM (Supplementary Fig. 2).

Isothermal calorimetry (ITC) was used to measure binding between Indisulam and the purified proteins in the absence of RBM39 (Fig. 1e). Indisulam binds the DCAF15-DDB1(ΔBPB)-DDA1 complex with a weak affinity of approximately 17 µM K_d_. This weak interaction was confirmed by ^1^H saturation transfer difference (STD) NMR, where STD peaks for Indisulam were only detected in the presence of 1 µM DCAF15-DDBA1-DDA1 (Supplementary Fig. 3). No STD peaks were detected between Indisulam and RBM39(Δ150) alone (indicating a K_d_ of >50 µM). Consistent with previous reports, Indisulam binds potently in the presence of both the DCAF15-DDB1(ΔBPB)-DDA1 complex and RBM39(Δ150) (187 nM K_d_), suggesting that Indisulam engages both DCAF15-DDB1-DDA1 and RBM39 to form a quaternary complex^17, 18^.

## Structure determination of DCAF15-DDB1-DDA1-Indisulam-RBM39 complexes by X-ray crystallography and cryo-electron microscopy

The three-dimensional structure of human DCAF15-DDB1(ΔBPB)-DDA1-RBM39(RRM2) in complex with Indisulam was determined by X-ray crystallography (Fig. 2). The human DCAF15-DDB1-DDA1-RBM39(RRM2)-Indisulam co-structure was solved by cryo-electron microscopy (Fig. 2c; Supplementary Fig. 4). The electron density maps for both structures were determined independently, thereby illustrating the structure by two separate methods. To allow determination of the most biologically relevant structure, care was taken to only minimally alter the proteins by mutation or deletion if necessary. The only necessary change was to delete the BPB domain of DDB1 to obtain large, well diffracting crystals for the X-ray studies; the sequence of DCAF15 was not modified. The resolution of the X-ray crystallography and EM structures are 2.3 Å and 3.5 Å, respectively. Crosslinking and mass spectrometer analysis of the DCAF15-DDB1-DDA1 complex provided important spatial constraints for the modeling and is described in Supplementary Fig. 5. Full statistics and methods are provided in Table 1 and the Methods section.

**Table 1.**
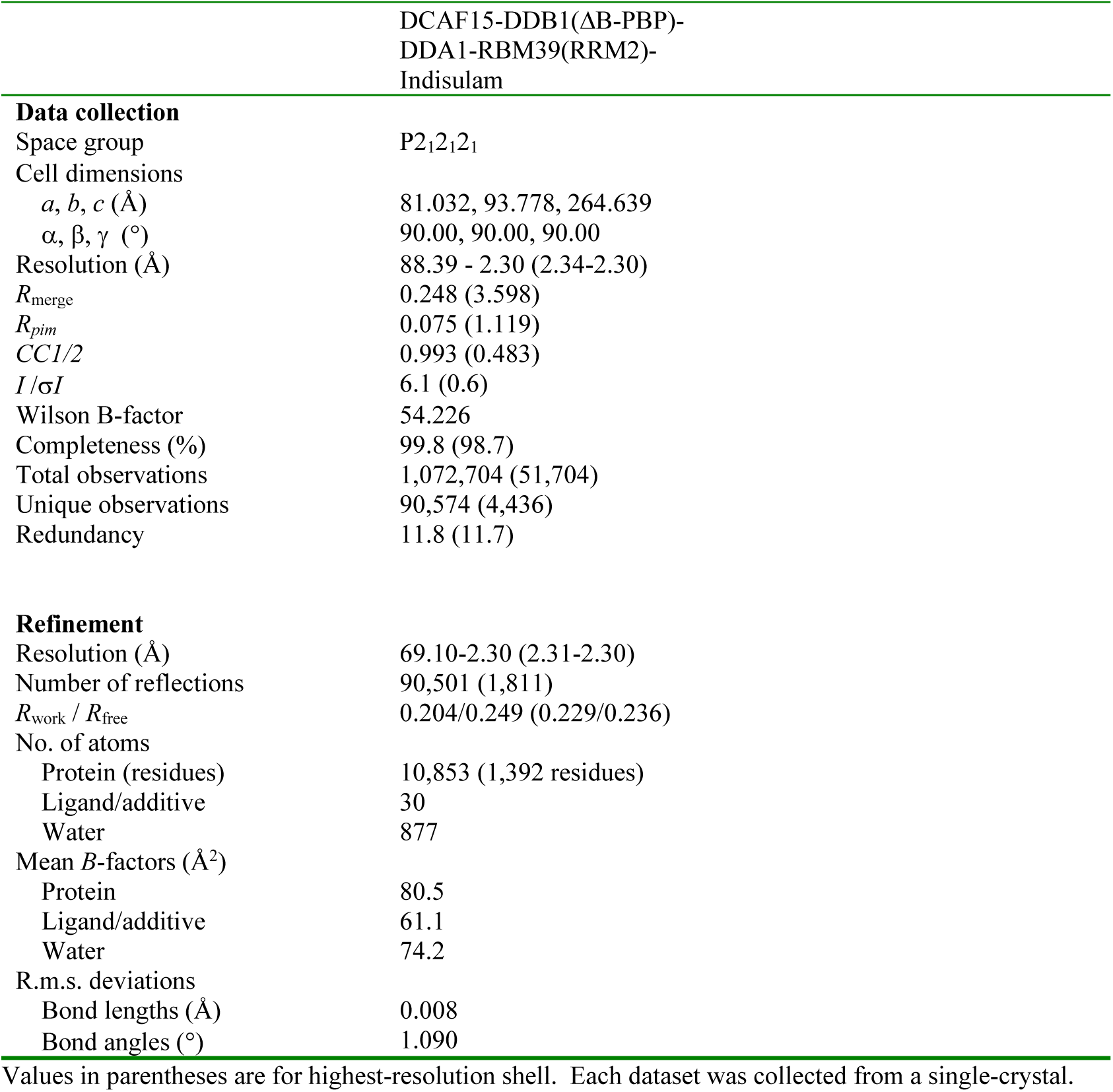
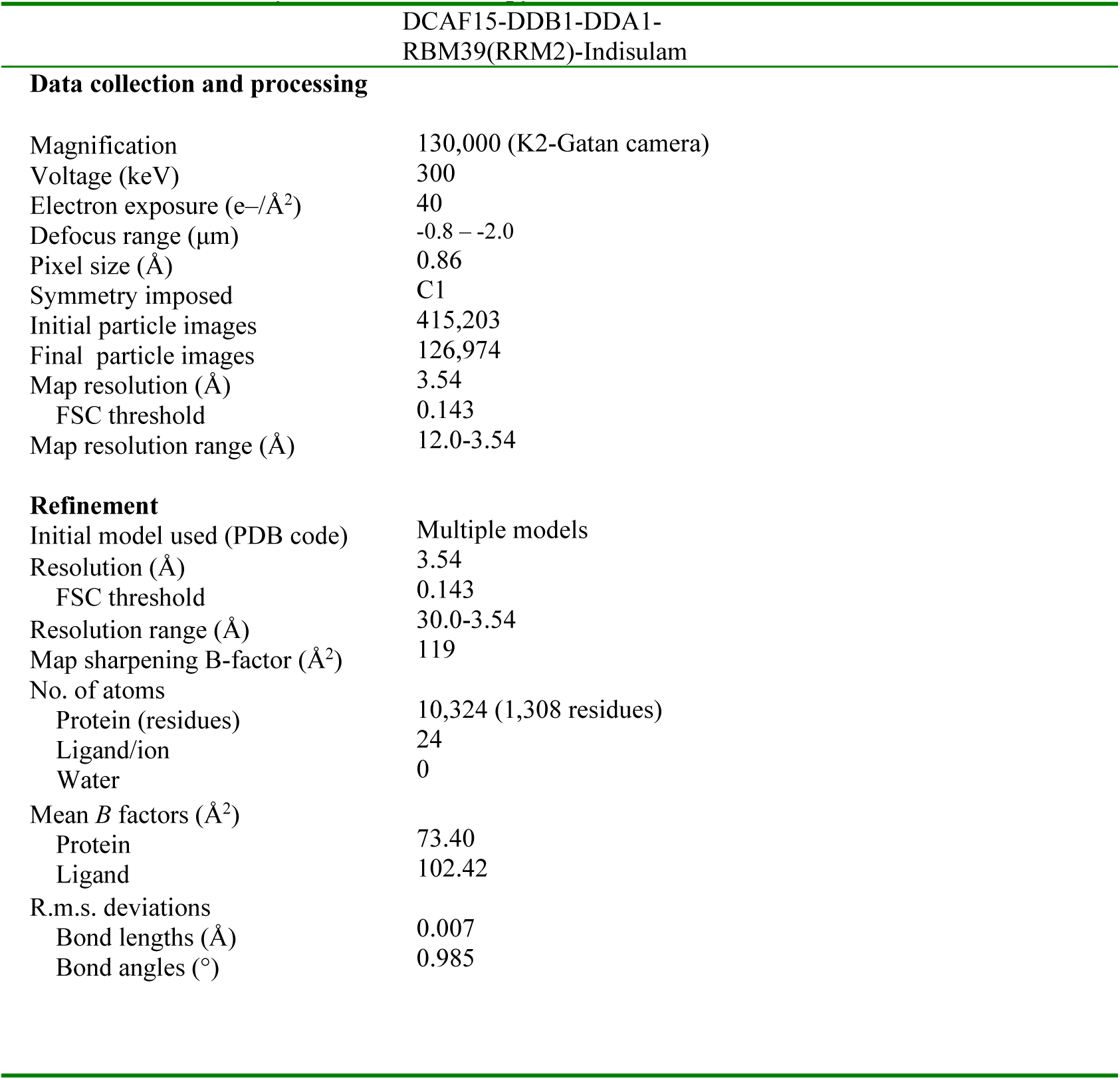
X-ray crystallographic data collection and refinement statistics (molecular replacement with iterative pseudo-atom phasing)

**Figure 2:**
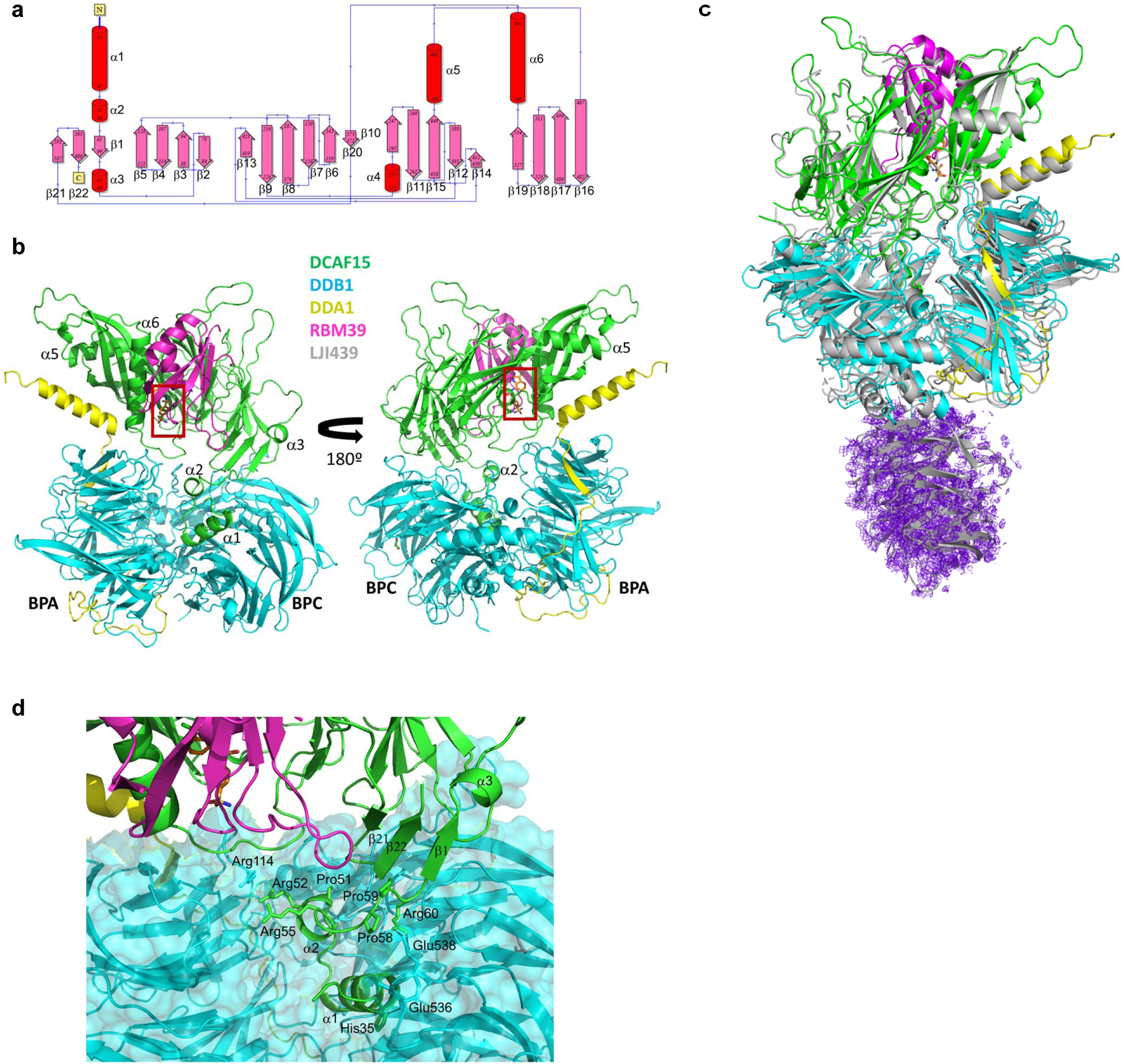
Structural analysis of the human DCAF15-DDB1-DDA1-RBM39(RRM2) complex with Indisulam. **a**, Secondary structure and connectivity diagram for DCAF15. Residues 1-31, 272-385, and 398-416 are disordered and are not visible in electron density maps. The N-and C-terminus are labelled. **b**, Overall quaternary structure of human DCAF15-DDB1(ΔBPB)-DDA1-RBM39(RRM2) in complex with Indisulam. DCAF15 is shown in green, DDB1 in blue, DDA1 in yellow, and RBM39(RRM2) in magenta. The Indisulam binding site between DCAF15 and RBM39(RRM2) is boxed in red. Two views separated by 180° are presented. Key structural elements on DCAF15 are labelled, as are the BPA and BPC domains on DDB1. **c**, The cryo-EM structure of human DCAF15-DDB1(ΔBPB)-DDA1-RBM39(RRM2) in complex with Indisulam overlapped with the X-ray co-structure. The cryo-EM co-structure is shown in grey. **d**, Helix-loop-helix docking interactions with DDB1 and the ‘arginine ladder’. The helix-loop-helix comprising a1 and a2 is shown docking to DDB1. Key hydrogen-bonding interactions are shown as dotted lines. Key hydrophobic residues are also shown. The unusual ‘arginine ladder’ comprised of Arg52 and Arg55 from DCAF15 and Arg114 from DDB1 is also shown, as are portions of DDA1, RBM39(RRM2), and Indisulam

## The structure and topology of DCAF15

DCAF15 forms direct interactions with DDB1, DDA1, and RBM39(RRM2) in the multi-protein complex (Fig. 2b). Full-length DCAF15 comprises a novel fold of 6 α-helices and 22 largely anti-parallel β-sheets (Fig. 2a)^19^. DALI analysis^20^ suggests some topological similarity with WD repeats from proteins such as WD40 repeat containing protein 5 (PDB accession code 4CY1^21^ and others^22, 23^) and SWD1-like protein (PDB accession code 6E29^24^), but the root-mean-square-deviation (RMSD) overlap with these proteins is quite high (>4 Å), indicating that there are only disparate regions of structural similarity. Moreover, the DCAF15 fold is topologically less symmetric than these domains which suggests that DCAF15’s fold is distinct from typical WD-type domains^25^. DCAF15 exhibits three disordered regions consistent with PONDR^26^ predictions: residues 1-31 at the N-terminus and residues 272-385 and 398-416 in the middle of the protein. The remainder of DCAF15 is well-ordered, including the C-terminus, which is sequestered within the body of the protein, proximal to the N-terminus.

Near the N-terminus of DCAF15 is a helix-loop-helix (residues 35-59) which mediates its interaction with DDB1, a feature shared with other CRL4 E3 ligase substrate receptors^27–29^ (Fig. 2d). The helix-loop-helix inserts into the large cleft formed between the BPA and BPC domains of DDB1. It is positioned into the cleft by a salt bridge between Arg60 of DCAF15 and Glu538 of the DDB1-BPC domain and interacts mainly with DDB1 by nonpolar shape complementarity with occasional side-chain mediated hydrogen bonds. The helix-loop-helix also contributes to an unusual feature of unknown significance, an ‘arginine ladder’, where Arg52 and Arg55 from the DCAF15 helix-loop-helix and Arg114 from the DDB1 PBA domain stack against each other and point to the same approximate region in space. The relative orientation of the two helices is ensured by complimentary hydrophobic packing on one side of each helix and the motif is ended by two consecutive prolines, Pro58 and Pro59.

## DDA1 stabilizes the DCAF15/DDB1 complex

DDA1 is highly-ordered: residues 4-44 form a strand which snakes around the surface of DDB1, residues 45-49 form a β-strand, and residues 53-76 form an α-helix (Fig. 3a). Residues 1-3 and 77-102 of DDA1 are disordered, consistent with PONDR^26^ predictions. The N-terminus of DDA1 binds to the DDB1-binding groove identified by Shabek and colleagues (PDB accession code 6DSZ)^30^. Interactions are mostly hydrophobic in nature, with insertion of aromatic groups into hydrophobic pockets a reoccurring theme (Tyr11, Phe16, and Phe19 on DDA1). The strand then continues along the face of DDB1, engaging hydrophobic pockets and forming main-chain hydrogen-bonding interactions until the start of the α-helix with residue 53. In its path over the surface of DDB1, DDA1 interacts with both the β-sheets and the loops between them. In many locations along this path, the hydrogen-bonding pattern of the main-chain to areas of DDB1 is equivalent to that of a parallel β-sheet.

**Figure 3:**
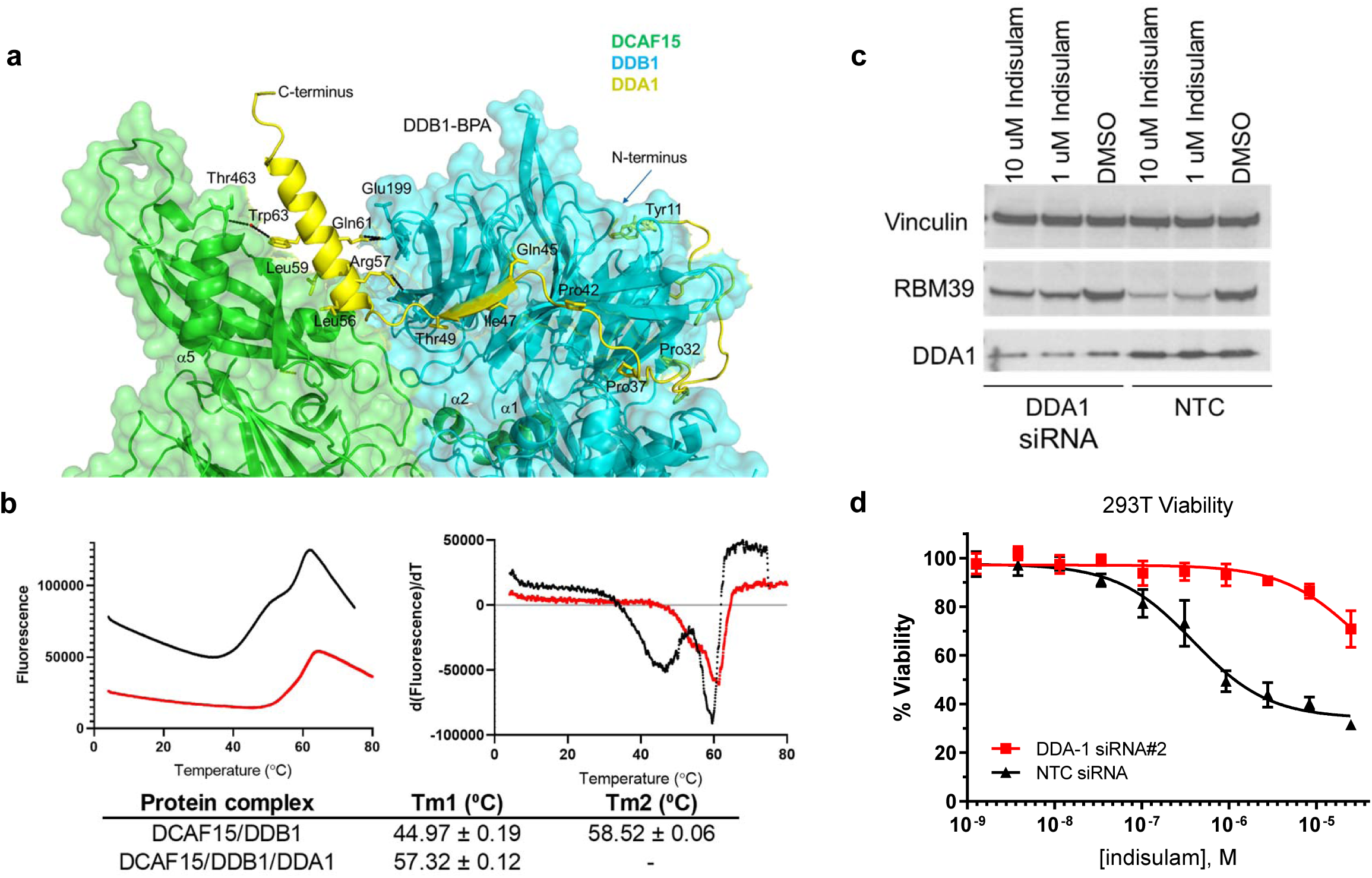
DDA1 stabilizes the DCAF15-DDB1 complex and impacts degradation of RBM39 by Indisulam. **a**, Interactions between DDA1(yellow), DCAF15 (green), and DDB1 (blue). Key residues are labelled and key salt-bridges and hydrogen-bonding interactions are shown as dotted lines. Residues on DDA1 which line the interaction surface for DDB1 are labelled. Within the β-sheet portion of DDA1, hydrogen-bond patterns similar to parallel β-sheet hydrogen-bonding exist. The N- and C-terminus of DDA1 are identified. **b**, Differential scanning fluorimetry (DSF) analysis measuring thermal stability of purified DCAF15-DDB1 (5 µM) (black lines) and DCAF15-DDB1-DDA1 (5 µM) (red lines) complexes. Both raw fluorescence (left) and – d(fluorescence)/d(temperature) (right) were plotted over temperature. Plotted data represent the median value for three (n=3) biological replicates from one individual experiment. Tabular Tm values are listed ± s.d. of the mean for the same three (n=3) biological replicates. **c**, Western blots showing levels of RBM39 in HEK293T cells transfected with DDA1 siRNA or a non-targeting control following 6 h treatment of 10 µM Indisulam, 1 µM Indisulam, or DMSO. Data shown from one individual, representative experiment from three independent repeats. **d**, Effect of 72 h Indisulam treatment on viability (CellTiterGlo) of HEK293T cells transfected with DDA1 siRNA (red line) or a non-targeting control (black line). Error bars represent s.d. of the mean from four biological replicates (n=4) in a single experiment. Each experiment was performed two independent times.

Residues 53-76 of DDA1 form an α-helix which serves to help anchor DCAF15 to DDB1 by bridging interactions between the two proteins. The face of the helix facing towards DCAF15 is predominately hydrophobic, with key polar residues forming specific interactions. For example, DDA1(Trp63) forms a structural water-mediated hydrogen-bond with main-chain amide of DCAF15 (Thr463) and DDA1(Lys66) forming a hydrogen-bond with the main-chain carbonyl of DCAF15(Val533). The opposite face is predominately hydrophilic and makes both direct and water-mediated interactions with the BPA domain of DDB1. For example, DDA1(Arg57) forms a salt-bridge with both the main-chain carbonyls of DDB1(Asn156) and DDB1(Lys200), while DDA1(Gln61) forms a hydrogen-bond with the main-chain carbonyl of DDB1(Glu199). Consequently, reconstitution and differential scanning fluorimetry (DSF) reveals greater stability of the DCAF15-DDB1-DDA1 complex compared to DCAF15-DDB1 alone (Fig. 3b). Moreover, knockdown of DDA1 in 293T cells impairs Indisulam-mediated degradation of RBM39 and the subsequent reduction in cell viability (Fig. 3c,d), confirming a functional role for DDA1 in DCAF15 cellular activity.

## RBM39-DCAF15 interactions observed in the complex

The RBM39(RRM2) domain has a typical RNA-recognition motif structure comprised of two α-helices positioned against four anti-parallel β-sheets^31, 32^. The RRM2 domain is positioned into a cleft existing between α6 and β20 of DCAF15 with the RRM2 central α-helix (residues 261-273) positioned proximal to β9 and α6 in DCAF15. While the majority of the interactions between RBM39(RRM2) and DCAF15 are nonpolar, the majority of the polar interactions occur between DCAF15 and the central alpha helix of RBM39 (Fig. 4a,b). RBM39(Glu271) positions RBM39(Arg267) via a salt-bridge to coordinate π-π interactions^33^ with DCAF15(Phe139) and DCAF15(Phe157) (Fig. 4a); RBM39(Glu271) also forms a direct salt-bridge with DCAF15(Arg178). These interactions are important for Indisulam activity, as the Glu271Gln mutation reduces RBM39 recruitment to DCAF15 by ∼1000-fold as measured by fluorescence polarization (Fig. 4c,d). RBM39(Pro272) is positioned within a small hydrophobic pocket on DCAF15 and, like RBM39(Gly268), maintains close surface contact between RBM39 and DCAF15. Disrupting these contacts by Gly268Val or Pro272Lys mutations ablates Indisulam-induced RBM39 binding. A Pro272Ser mutation is better tolerated, but lowers binding affinity by ∼6-fold, suggesting the importance of hydrophobic character at this position.

**Figure 4:**
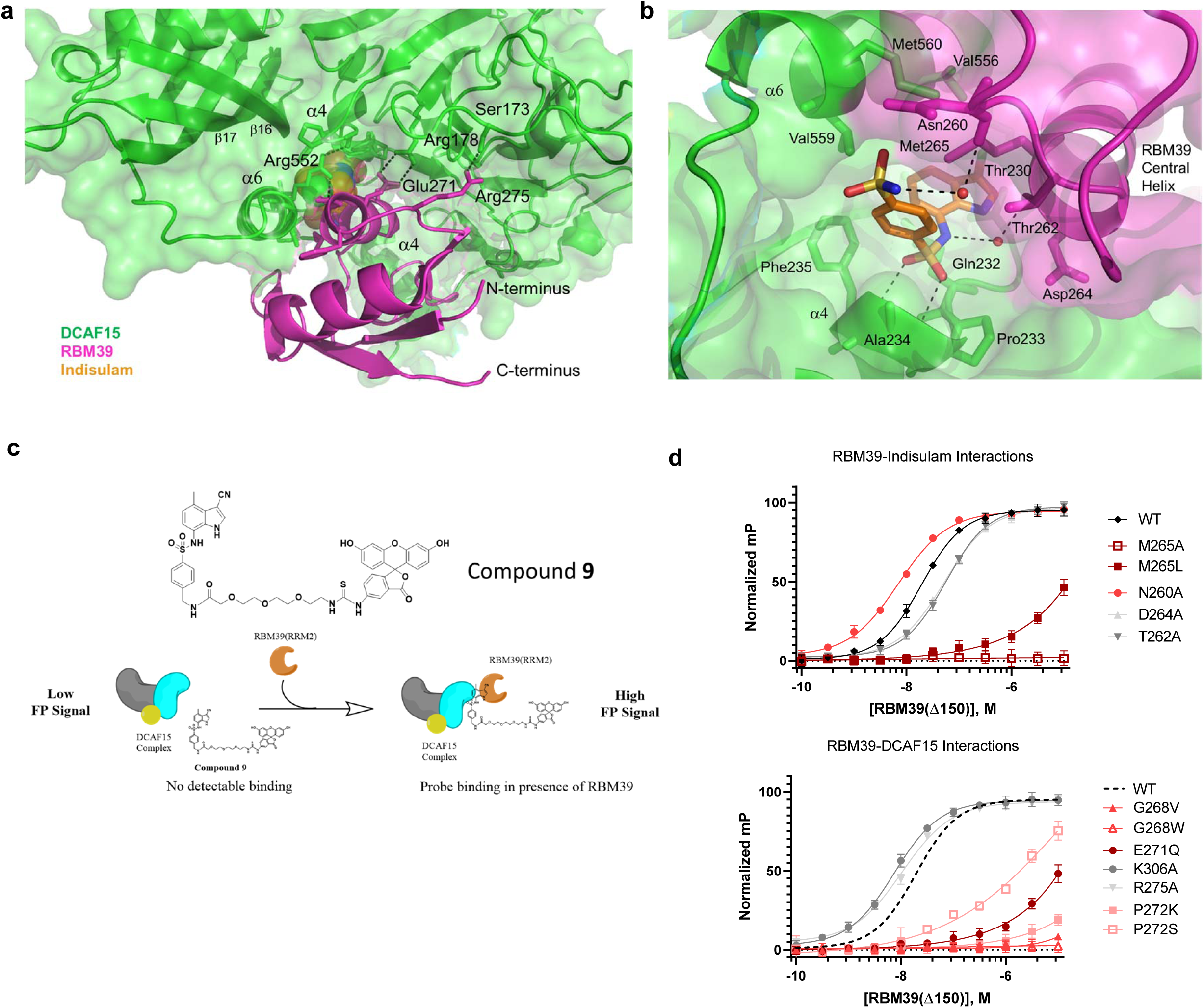
Detailed description of Indisulam binding at the DCAF15 and RBM39 interface. **a**, Non-Indisulam mediated interactions between DCAF15 (green) and RBM39 (magenta). Enthalpic interactions are shown as dotted lines. The majority of the interaction is due to shape complementarity and non-polar interactions, with interspersed electrostatic interactions and hydrogen-bonding. The N- and C-terminus of RBM39 are identified. **b**, Indisulam-mediated interactions between DCAF15 and RBM39. Indisulam (orange) bridges the structure of DCAF15 (green) and RBM39 (magenta) by forming several direct or water-mediated interactions with both DCAF15 and RBM39 and serves to increase the complementarity between the two surfaces. Hydrogen-bonds are shown as dotted lines and the surfaces of both DCAF15 and RBM39 are shown. Key residues are labelled. **c,** Schematic illustrating FP assay used to measure ternary complex formation between DCAF15-DDB1-DDA1, RBM39(Δ150) variants, and a FITC-labeled Indisulam analog **9**. 100 nM of DCAF15 complex is insufficient to bind **9** in the absence of RBM39(Δ150) and generates a low FP signal. In the presence of increasing concentrations of RBM39(Δ150) a ternary complex forms and FP signal increases. Protein titration data is described in Supplementary Fig. 9. **d**, FP assay measuring ternary complex formation between DCAF15-DDB1-DDA1 and RBM39(Δ150) variants bearing mutations at residues mediating direct and water-mediated interactions with Indisulam (top) or DCAF15 (bottom). Error bars represent s.d. of the mean from eight biological replicates (n=8) in a single experiment. Dotted line in right graph represents nonlinear data fit from wild-type RBM39(d150) data shown in left graph. Each experiment was repeated three independent times. Characterization data for all RBM39 variants are included in Supplementary Fig. 10

As nonpolar surface contacts are key contributors to RBM39-DCAF15 interaction, the MOE “patch analyzer” tool^34^ was used to characterize the RBM39(RRM2)-DCAF15 interface. Approximately 5.5% of the DCAF15 surface and 26.3% of the RBM39 surface is sequestered from solvent and engaged in protein-protein or protein-compound interactions. The largest hydrophobic patch (140 Å^2^) on RBM39 is formed by residues in and around the central helix and overlaps partially with a large hydrophobic patch present on DCAF15. In addition, there are four other hydrophobic patches in DCAF15 that are in contact with RBM39 and sequestered from solvent. The total nonpolar area on DCAF15 involved in the interaction with RBM39 is approximately 590 Å^2^, likely comprising the bulk of DCAF15-RBM39 binding energy. While non-polar interactions dominate the DCAF15-RBM39 interface, there are also occasional polar interactions at the periphery. For example RBM39(Arg275) hydrogen-bonds with DCAF15(Ser173), albeit with suboptimal geometry, while RBM39(Lys306) forms a weak hydrogen-bond with DCAF15(Thr543) (Fig. 4a). Interestingly, neither of these peripheral interactions appear critical for RBM39 recruitment, as substituting alanine for RBM39(Lys306) or RBM39(Arg275) is largely tolerated. Overall, while these DCAF15-RBM39 interactions are incapable of maintaining DCAF15 and RBM39 binding on their own (Fig. 1d), these largely nonpolar contacts are needed for Indisulam-mediated recruitment.

## The structural basis by which Indisulam enhances RBM39 binding to DCAF15

Indisulam binds between the RBM39(RRM2) central helix and β9, β16, and α6 of DCAF15 (Fig 4b). A polar cation-π interaction likely occurs between the heterocycle and DCAF15(Gln232), which is positioned adjacent to the chloro-indole group of Indisulam. The chloro-indole group of Indisulam binds in a hydrophobic pocket comprised of the aliphatic face of Thr230, Phe235 and Val559 from DCAF15, and also RBM39(Met265). RBM39(Gly268) also forms a periphery of the hydrophobic pocket. The phenyl-sulfonamide is positioned between several aliphatic sidechains, including DCAF15(Ala234), DCAF15(Thr262), and RBM39(Met265). Mutations at RBM39(Met265) and RBM39(Gly268) dramatically impact recruitment (Fig. 4d), suggesting that these residues contribute significantly to the DCAF15 binding-pocket.

The central sulfonamide accepts two hydrogen-bonds from the main chain amides of DCAF15 Ala234 and Phe235; the geometry of these interactions is near-optimal. It should be noted that both MoKa^35^ calculations and experimental pKa determination show that the nitrogen of the central sulfonamide bears a negative charge (Supplementary Fig. 6). This nitrogen forms water-mediated hydrogen-bonds with both RBM39(Thr262) and RBM39(Asp264). Given that alanine substitution at these positions abrogate binding by ∼2 fold, these interactions appear to contribute to RBM39 recruitment albeit modestly. The distal sulfonamide donates a hydrogen-bond from the nitrogen to a structural water which in turn hydrogen-bonds to the main-chain carbonyl of Asn260 of RBM39. The edge of this group exits the body of the complex towards bulk solvent. Altering the flexibility at this position by alanine substitution improves binding by ∼3-fold, illustrating its positive contribution.

From a conceptual view, Indisulam interactions largely complete the full complementarity lacking at the DCAF15-RBM39(RRM2) interface. While several polar interactions are found between Indisulam and RBM39, few appear crucial for recruitment. The most consequential perturbations involve disruption of nonpolar interactions with the indole and terminal phenyl group of Indisulam, as well as those that disrupt nonpolar DCAF15-RBM39 surface contacts.

## Using Indisulam analogs to identify important chemical features driving RBM39 recruitment to DCAF15

To assess the structural model and to better understand the plasticity of the small molecule binding pocket, Indisulam analogs were tested for their ability to recruit RBM39 to the DCAF15-DDB1-DDA1 complex using a biochemical time-resolved fluorescence resonance energy transfer (TR-FRET) assay (Fig. 5a). The chloro group at position R1 in Indisulam can be substituted by nonpolar groups of similar volume, such as methyl and nitrile (compounds **1**-**2**; Fig. 5b,c). The nitrile group, a known chlorine isostere, is especially effective and improves the EC50 to 1.21 µM. Substitution of the chloro group with a proton at position R1 (compound **3**) is not tolerated. The proton substitution removes ∼38 Å^2^ of hydrophobic surface interaction (∼0.95 kcal/mol of binding energy^33^) and would likely compromise the positioning of the remainder of the compound-protein contacts.

**Figure 5:**
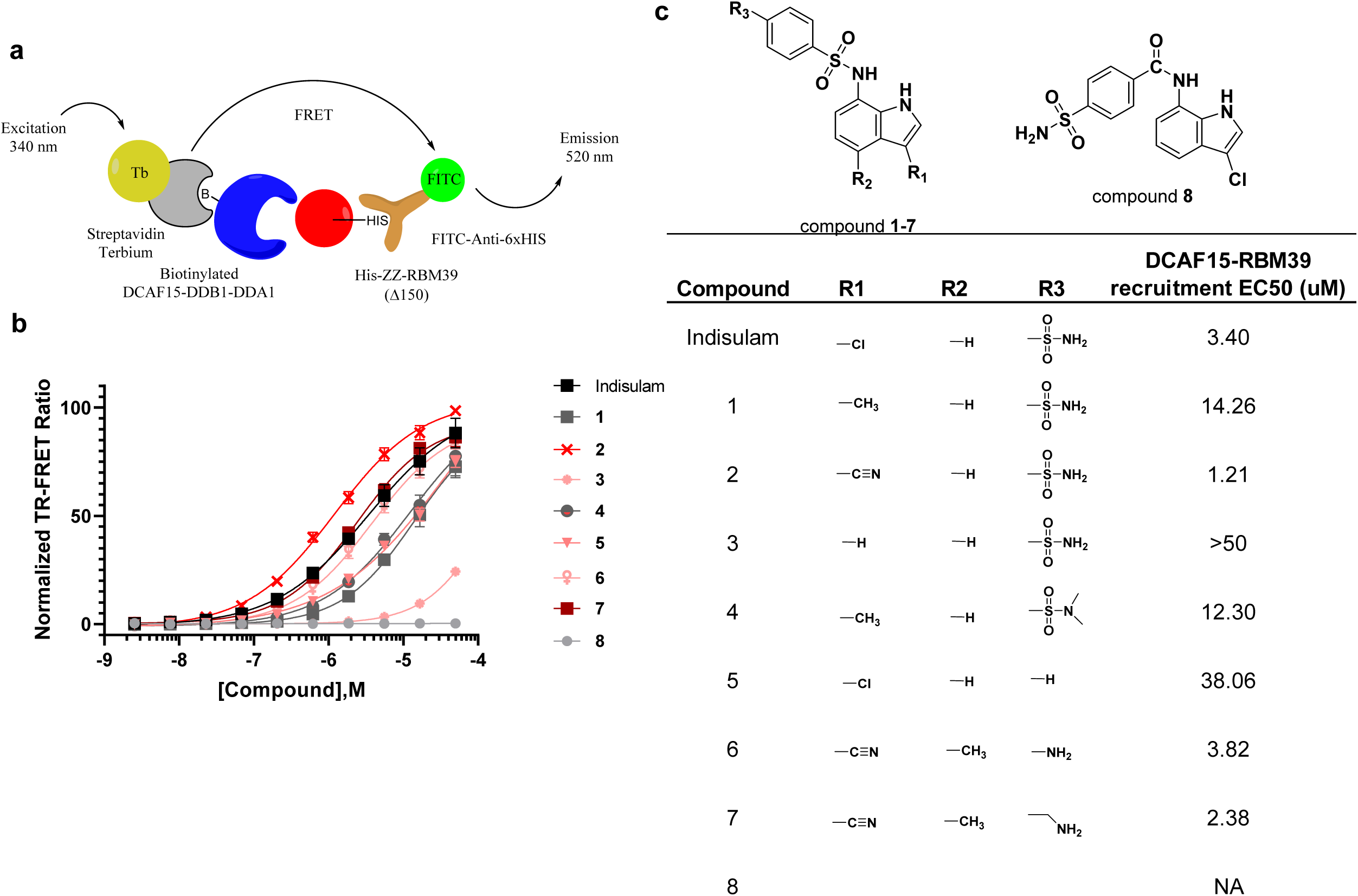
Structure-activity relationships for Indisulam as measured by DCAF15-DDB1-DDA1-RBM39 recruitment assay. **a**, Schematic illustrating time-resolved fluorescence energy transfer (TR-FRET) assay used to measure compound-induced recruitment of RBM39(Δ150) to DCAF15-DDB1-DDA1. Biotinylated DCAF15-DDB1-DDA1 complex was labeled with streptavidin-terbium conjugate to act as fluorescent donor to a FITC-antibody conjugate bound to his-tagged RBM39(Δ150). Upon ternary complex formation, TR-FRET signal is measured and reported as a ratio between emissions at 340 nm and 520 nM. **b**, DMSO-normalized TR-FRET ratios measured for varied doses of Indisulam (black line) and analogs in DCAF15-DDB1-DDA1-RBM39 recruitment assay. Error bars represent s.d. of the mean from four biological replicates (n=4) in a single experiment. **c**, Structure-activity relationship table describing generalized structures for analogs included (top) and impact of various substituents on DCAF15-DDB1-DDA1-RBM39 recruitment as described by EC_50_ values in TR-FRET assay (bottom).

The contributions of the terminal sulfonamide at position R3 are explored using compounds **4**-**7**. Compound **4** replaces the terminal sulfonamide with a dimethyl-sulfonamide at position R3 combined with a methyl replacement at position R1. This weakens the EC50 approximately 4-fold relative to Indisulam, roughly equivalent to the 4-fold difference for the methyl substitution alone. This suggests that the dimethyl-sulfonamide substitution is well tolerated. The two methyl groups can make hydrophobic contacts, maintaining non-polar surface area, while the nitrogen maintains hydrogen-bonding to the structural water which bridges its interaction to the backbone carbonyl of Asn260 on RBM39. In compound **5**, the terminal sulfonamide was replaced with a proton, removing the hydrogen bond donor at this position. RBM39 recruitment EC50 was reduced 11 fold suggesting a loss of ∼1.4 kcal/mol of binding energy which is consistent with the binding energy provided by a hydrogen-bond. In compounds **6** and **7** the terminal sulfonamide was replaced with an amine or methyl amine which are hydrogen bond donors. Additionally, the chloro group at position R1 was replaced with the effective nitrile and a methyl group replaced the hydrogen at position R2. Compounds **6** and **7** maintain strong RBM39 recruitment suggesting that the terminal sulfonamide can be replaced by other hydrogen bond donor groups.

Compound **7** degrades cellular RBM39 and reduced HCT116 viability with similar potency to Indisulam (Supplementary Fig. 7). STD NMR epitope mapping predicts that the nitrile-methyl-indole ring of compound **7** interacts with DCAF15, whereas the central phenyl group is more exposed to solvent in the absence of RBM39 (Supplementary Fig. 8). These data are consistent with the predicted binding pose of Indisulam and suggest that the terminal amine on **7** may represent an ‘exit vector’ for attachment of large substituents. We have confirmed these findings by solving the DCAF15-DDB1(ΔBPB)-DDA1-RBM39(RRM2) X-ray crystal structure in complex with **7** (data not shown), which shows that its binding pose is equivalent to Indisulam’s.

Lastly, Compound **8** probes the importance of the central sulfonamide in the recruitment of RBM39. Replacement of the central sulfonamide with an amide ablates RBM39 recruitment. This modification disrupts hydrogen-bonding with the backbone amides on DCAF15, and the amide linker also significantly alters conformational preferences of compound 8, thereby disrupting its binding. Overall, the observed compound SAR supports the predicted binding mode for Indisulam and highlights regions amenable to further modification.

## Indisulam selectivity predictions based on critical RBM39 residues

A question of great interest is whether new, yet unidentified proteins can be recruited to DCAF15 by Indisulam. The structure of the complex, RBM39 mutagenesis studies and previously published work^17, 18^ suggest that RBM39 residues Met265, Gly268, Glu271, and Pro272 in the central alpha helix are necessary for Indisulam-mediated DCAF15 binding. An alpha helical “X^1^XXM^4^XXG^7^XXEP^11^” sequence was defined as a putative degron motif.

The bioinformatics workflow is summarized in Figure 6a. To identify compatible proteins with the putative degron motif, 20,421 unique human protein sequences from uniprot were accessed (http://www.uniprot.org). 6,475 of these entries had associated x-ray/NMR structures. For proteins with known structures, 3,425 proteins were identified with a glycine at position 7 of the alpha helix, and a helix RMSD of less than 2Å. The helix RMSD is determined by first aligning all identified helices containing glycine at the correct position to the RBM39(Thr262-Pro272) central helix from the DCAF15 co-structure, and then calculating a backbone RMSD of the two helices using the alpha-carbons. Next, steric clashes of these target proteins with DCAF15 were calculated, where clashes of less than 10 heavy atoms between DCAF15 and the target protein were considered acceptable. This yielded 1787 targets with good helix overlay and minimal steric clash. While these protein entries may be sterically compatible with Indisulam-bound DCAF15, only the helices from RBM39 and RBM23 comprised sequences matching our putative degron motif. Expression proteomics studies in HCT116 cells treated with 10 µM indisulam for 4 hours showed that RBM39 and RBM23 were the most significantly downregulated proteins (>2 fold over DMSO, Fig. 6b). While ZNF277 was also identified as a potential target from these studies, Western blot analysis confirmed that only RBM23 and RBM39 were reduced by indisulam (Fig. 6c).

**Figure 6:**
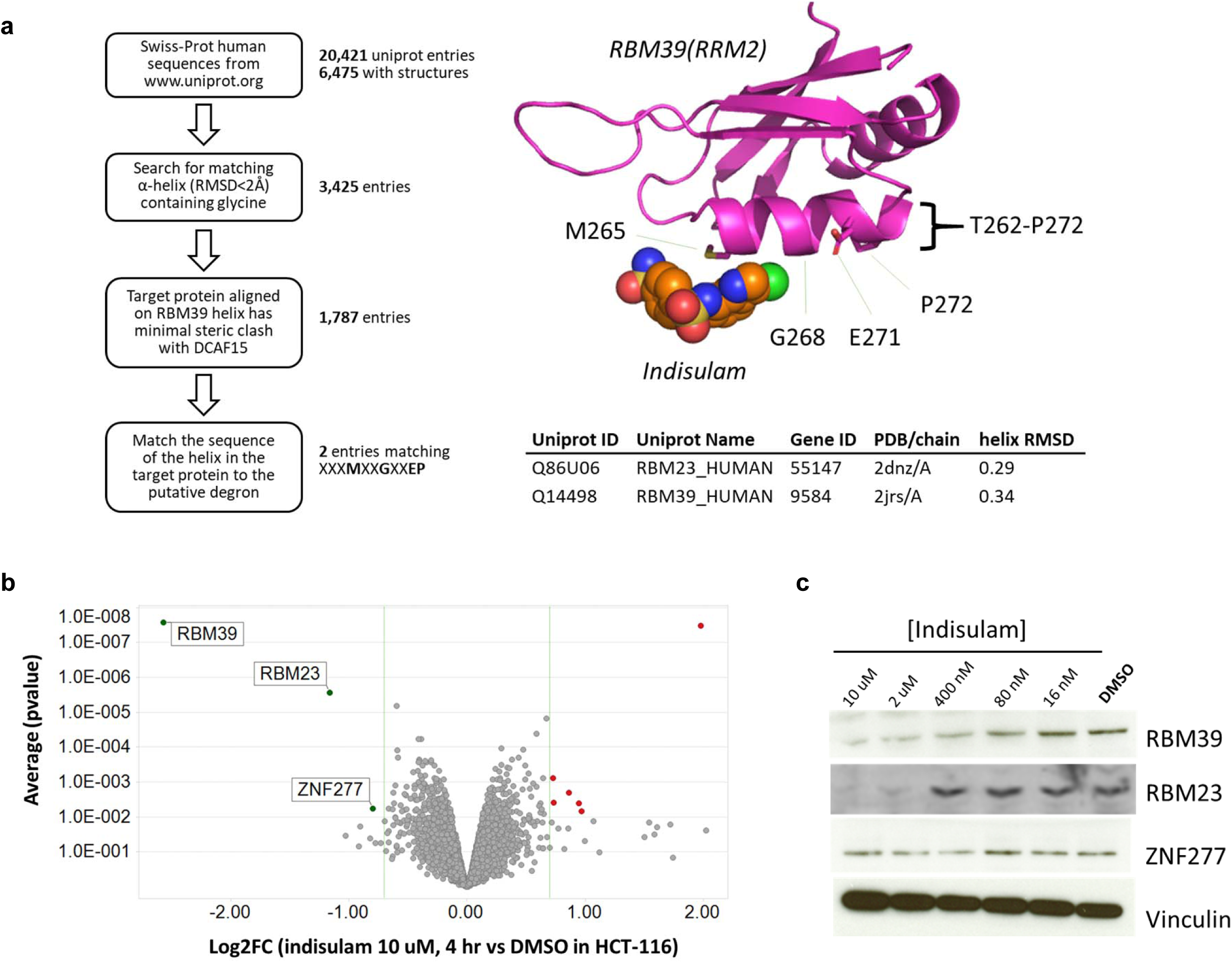
Proteome-wide motif search predicts Indisulam selectivity confirmed by expression proteomics. **a**, Bioinformatics workflow for identifying proteins bearing putative degron motif required for DCAF15-Indisulam recruitment (left) and structures of DCAF15-bound Indisulam (orange) and RBM39(RRM2) (magenta) with central alpha helix highlighted (right). The RBM39 residues found to be most critical for DCAF15-Indisulam recruitment are labeled (M265, G268, E271, and P272). 6,475 proteins with known structures were identified in the Swiss-Prot database, of which 3,425 had a glycine in an alpha helix. 3,112 of the glycine-containing alpha helices aligned to an RBM39(RRM2) structure (2JRS) with RMSD <2.0 Å. Among these matches, only RBM23 and RBM39 helices had a sequence matching the required X^1^XXM^4^XXG^7^XXEP^11^ motif. RMSD values, PDB IDs, and gene names shown in bottom table. **b**, Volcano plot summary of expression proteomics experiments comparing lysates from HCT-116 cells treated for 4 h with 10 µM Indisulam or DMSO. Significant downregulated proteins (p value < 1E-2, Log_2_ fold-change) are labeled. Data represents two (n=2) biological replicates per treatment condition in a single experiment. **c**, Western blots showing levels of RBM39, RBM23, and ZNF277 in HCT116 cells following 4 h treatment with varied concentrations of Indisulam or DMSO. Data shown from one individual, representative experiment from three independent repeats.

## Discussion

TPD is a promising new area of drug discovery focused on developing small molecules that bring “difficult-to-drug” targets in close proximity to E3 ligases to induce UPS-dependent target degradation^12^. Understanding the mechanism of action of the few known TPD molecules will drive progress in this young field. Our work leveraged structural biology, biophysics, and chemical and genetic variomics to study the TPD molecule Indisulam and to understand precisely how Indisulam recruits RBM39 to the DCAF15 E3 ligase substrate receptor.

The co-structure reveals that Indisulam behaves as a molecular glue degrader rather than an allosteric inducer of dimerization like Gibberellin^12^. DCAF15, with its novel fold, embraces RBM39(RRM2) with Indisulam interacting with both proteins to promote a suitable interface (Fig. 2b). Indisulam sits in a well-defined pocket formed by DCAF15 and coordinates several direct and water-mediated interactions with both RBM39 and DCAF15 (Fig. 4b). There is also a series of interactions between DCAF15 and RBM39. These DCAF15-RBM39 interactions are insufficient to enable binding on their own, as interaction in the absence of Indisulam could not be detected by AUC (Supplementary Fig. 2) or SPR (data not shown). It is therefore unlikely that Indisulam binds to a site pre-formed by RBM39 and DCAF15 association or stabilizes a basal interaction, ruling out a brefeldin A-like mechanism^36^. Given that Indisulam fails to bind RBM39 alone (Fig. 1e), it is not a direct destabilizer of RBM39, like the SERDs^10^. Indisulam does bind the DCAF15-DDB1-DDA1 complex alone with weak affinity (∼17 uM, Fig. 1e) and binds more potently in the presence of RBM39 (187 nM, Fig. 1e). This ∼100-fold binding affinity enhancement likely stems from the additional direct and water-mediated polar contacts made between RBM39 and Indisulam, along with the series of nonpolar interactions emerging from the apparent structural complementarity between DCAF15 and RBM39. Overall, the data suggests that an Indisulam-DCAF15 interaction precedes association with RBM39, whose recruitment in turn stabilizes Indisulam binding.

This report provides the first high-resolution structural view of the DCAF15 E3 ligase, a DDB1-CUL4 associated substrate receptor about which little is known. Apart from the reported association with its partners DDA1, DDB1, and associated CRL4 components^17, 18^, no additional endogenous binding partners nor substrates have been identified to date. Human papillomavirus (HPV) E6 and E7 proteins, known oncogenic factors capable of recruiting host E3 ligases for the degradation of tumor suppressors^37, 38^, were found to bind DCAF15, however, the biological consequences of this event are unclear^39^. Given that DCAF15 exhibits significant disorder in our co-structure and comprises a novel fold distinct from the WD40-type domain seen among other CRL4 E3 substrate receptors (e.g. CRBN, DDB2), additional binding partners unique to DCAF15 are likely to be implicated in its biology and may help to facilitate the identification of novel substrates. Like other DDB1-associated E3 ligases, a helix-loop-helix mediates interaction between DCAF15 and DDB1 along with several hydrogen-bonding, water-mediated, and nonpolar interactions (Fig. 2d).

DDA1, a member of the E2-interacting DDD complex^40^ with de-etioloated 1 (DET1) and DDB1, was pulled down with DCAF15 in the presence of Indisulam^17, 18^. While immunoprecipitation (IP) studies have found DDA1 to be associated with several CRL4-DCAFs with the exception of DDB2^41^, its impact on CRL4 biology is largely unknown. A recent report suggests that DDA1 regulates lenalidomide-mediated ubiquitination of IKZF1/3 through its association with CRBN-DDB1^42^, although CRBN IP studies fail to capture DDA1 as a binding partner^43, 44^. DDA1 improves the thermal stability of the DCAF15-DDB1 complex and is required for Indisulam-mediated degradation of RBM39 (Fig. 3), suggesting a significant role for DDA1 in DCAF15-CRL4 complexes. The N-terminus of DDA1 binds to a previously reported DDB1-binding groove^30^, while the C-terminus forms an α-helix which serves as an interface to help anchor DCAF15 to DDB1 through bridging interactions. Crosslinking mass spectrometry studies (Supplemental Fig. 6) suggest DDA1 displays dynamic mobility, signifying possible roles in substrate recognition or ubiquitination. Further work will be required to elucidate the potential roles of DDA1 in CRL4 ubiquitination.

A key question for drug discovery and design is whether Indisulam and related aryl sulfonamides analogs could potentially provide a route to novel DCAF15-based molecular glue degraders, parallel to the development of IMiD analogs capable of degrading a diversity of targets through CRBN^14^. From a compound perspective, IMiDs bind CRBN with high affinity (100-200 nM)^14, 29^ whereas Indisulam binds DCAF15 weakly (∼17 µM). For Indisulam, RBM39 target binding is needed to potently engage the DCAF15 E3 ligase. Regions of Indisulam required for DCAF15 binding overlap with regions needed for target recruitment (Fig. 4b), unlike IMiDs where there is a separation between the CRBN binding region, the glutaramide warhead, and the recruitment region. IMiD-CRBN complexes appear uniquely poised to recruit proteins bearing a beta hairpin structural motif^14, 45^. This is achieved primarily through IMiD-bound CRBN interactions with a precise spatial arrangement of backbone on the beta hairpin degron^29, 45–47^. Selectivity for a given beta hairpin protein can be achieved through compound-mediated interactions with unique side-chain features on the recruited target^14, 29, 46, 47^. DCAF15-bound Indisulam primarily engages with side-chains of the RBM39 target and coordinates only a few interactions with backbone elements (Fig. 4b), likely underpinning its remarkable selectivity. Proteomics analysis of HCT116 cells treated with 10 µM Indisulam for 4 hours only showed downregulation of RBM39 and RBM23 (Fig. 6b,c), two RRM domain-containing protein sharing nearly 89% identity within these RRM domains. As our bioinformatics analysis reveals, only these two proteins harbor the requisite glycine-containing alpha helix with critical residues able to mediate interactions with both Indisulam and DCAF15 (Fig. 6a). Moreover, RBM23 harbors this sequence within an RRM domain, likely enabling further nonpolar interactions via complementarity with DCAF15. While other RRM-domain proteins may complement the DCAF15 binding pocket, none bear the critical motif necessary to engage Indisulam to a degree similar to RBM23 or RBM39. We propose that novel DCAF15-binding chemotypes would be required to engage DCAF15 and recruit additional partners in a programmable manner similar to IMiD analogs. DCAF15 binders that separate ligase binding and target recruitment and which coordinate backbone features of a recruited degron could potentially provide a route to programmable DCAF15 degraders. However, such compounds and structural degrons have yet to be identified and this represents the challenge for the development of future DCAF15 molecular glues degraders.

## Methods

### Cloning, protein expression and purification

The gene for DCAF15 was codon-optimized and cloned into a pFastBac vector with an N-terminal His-ZZ-3C tag. DDB1, DDB1-ΔBPB (DDB1 with a residue 398-701 deletion), and DDA1 were each cloned into a pFastBac vector without an associated tag. The pFastBac constructs were used to generate baculovirus using the Bac-to-Bac method (Bac-to-Bac® Baculovirus Expression System, ThermoFisher). The baculovirus was amplified in SF21cells (ThermoFisher). DCAF15 was co-expressed with DDB1 (or DDB1-ΔBPB) and DDA1 in SF21 cells seeded at a density of 1.5 x 10^6^ cells/mL and synchronously infected with the recombinant baculoviruses at a volume of 2%:4%:4%. The cultures were grown in 2-L glass Erlenmeyer flasks in serum-free media with agitation at 120 rpm for 48 hours at 27 °C. Cells were harvested two days post-infection, flash frozen, and stored at −80 °C.

Cell pellets were resuspended in lysis buffer consisting of 50mM HEPES, pH 7.5, 500 mM NaCl, 20mM imidazole, 10% Glycerol (v:v), 2mM TCEP, universal nuclease (Pierce), and 3x protease inhibitor (Roche) and lysed using a Dounce homogenizer. The clarified cell lysate was mixed with 10 mL Ni-NTA resin and incubated at 4 °C for 4 hours. The resin was washed with IMAC buffer A, which consisted of 50mM HEPES, pH7.5, 500 mM NaCl, 20mM Imidazole, 10% glycerol (v:v), and 2mM TCEP. The protein was eluted with IMAC buffer B, which consisted of 50mM HEPES, pH7.5, 500 mM NaCl, 500mM imidazole, 10% glycerol (v:v), and 2mM TCEP. The fractions containing the His-ZZ-3C-DCAF15-DDB1-DDA1 complex were combined and treated with 3C protease overnight in dialysis buffer consisting of 50mM HEPES, pH7.5, 400 mM NaCl, and 1mM TCEP. The cleaved protein was then purified by a 5 ml HisTrap HP column on an AKTA Avant system (GE Healthcare). The flow through was combined and diluted with buffer C, which consisted of 50mM HEPES pH7.5, and 2mM TCEP, and purified by a 5 mL HiTrap Q HP column. The protein was eluted with a 100 mL linear gradient of 200 mM to 500 mM NaCl in buffer C. Fractions containing the complex were concentrated and further purified by a Superdex 200 26/60 column (GE Healthcare) in buffer D, which consisted of 50mM HEPES, pH7.5, 300 mM NaCl, and 1mM TCEP. The yield of purified complex following this procedure was approximately 15 mg of DCAF15-DDB1-DDA1 per liter of culture. The molecular weights of the proteins in the complex were confirmed by LC/MS. The intact protein mass was detected by LC/MS on an Open Access MS system (Agilent 1290 UHPLC + Agilent 6530 QToF) and analyzed by the MassHunter software.

RBM39 (residues 151-end) and RBM39 (residues 250-328) were each cloned into pET30b vector with an N-terminal His-ZZ-3C tag. Q5 mutagenesis (NEB) was performed on these constructs to generate reported mutants using manufacturer’s protocol. Proteins were expressed in *E. coli* BL21(DE3) cells (16 ⁰C for 18 hrs followed byinduction by 1 mM IPTG). Cell pellets were resuspended in lysis buffer containing 50mM HEPES, pH 7.5, 500 mM NaCl, 20mM imidazole, 10% Glycerol (v:v), 2mM TCEP, and 1x HALT protease inhibitor cocktail (ThermoFisher). After lysis by sonication, the protein was purified by a HisTrap column, cleaved by 3C protease, and purified again by a HisTrap column. The cleaved protein was further purified by a Superdex 200 16/60 column (GE Healthcare) in 50mM HEPES, pH7.5, 300 mM NaCl, and 1mM TCEP. For His-ZZ-RBM39(151-end) variants, proteins were purified by batch Ni-NTA bead purification (1 mL slurry/1L culture; Qiagen) and further purified by a Superdex 200 16/60 column (GE Healthcare) in 50mM HEPES, pH7.5, 300 mM NaCl, and 1mM TCEP.

### Surface plasmon resonance binding analysis

500 resonance units (RU) of Biotinylated DCAF15-DDB1-DDA1 was immobilized to a streptavidin SA Sensor Chip (GE Healthcare Life Sciences) in a running buffer consisting of 50 mM HEPES, pH 7.5, 300 mM NaCl, 1 mM TCEP, 0.05% (v:v) Tween-

20 supplemented with 20 μM Indisulam. All SPR studies were performed on a Biacore T200 instrument (GE Healthcare Life Sciences). Following capture, any remaining streptavidin sites on both the reference and active channels were blocked with biocytin. The association and dissociation steps of a dilution series of RBM39 (Δ150) or RBM39(RRM2) domains were performed in running buffer in the presence of excess Indisulam. All sensorgrams shown are reference subtracted with solvent correction procedures implemented. Data were fit to both steady state and a 1:1 kinetic model using Biacore evaluation software and gave equivalent dissociation constants (Kd).

### Analytical ultracentrifugation

Experiments were performed in a Beckman Optima analytical ultracentrifuge equipped with double sector, charcoal-filed centerpieces (12 mm path length, sapphire windows). In brief, 2.5 µM DCAF15-DDB1(ΔBPB)-DDA1 and 10 µM of His-ZZ-RBM39(Δ150) were incubated with and without 12.5 uM Indisulam for 60 min in 50 mM HEPES, pH 7.5, 300 mM NaCl, and 1 mM TCEP and subjected to sedimentation velocity at 42,000 rpm for 5 h at 20 °C. For analysis, buffer density, viscosity, and partial specific volumes (derived from amino acid composition) were calculated using SEDNTERP^48^. Rayleigh interferometric fringe displacement sedimentation data was collected and modeled with diffusion-deconvoluted sedimentation coefficient distributions c(s) in SEDFIT^49^.

### Isothermal calorimetry

Isothermal calorimetry was used to measure the heat of enthalpy of the DCAF15-DDB1-DDA1-RBM39(Δ150)-Indisulam complex formation using a GE Healthcare autoITC200 at 25 °C. Protein solution in the calorimetric cell containing both DCAF15-DDB1(ΔBPB)-DDA1 complex at 10 uM and RBM39(Δ150) at 10 uM were titrated with Indisulam at 100 uM, in carefully matched buffers. 19 injections were carried out until the proteins were fully saturated. The control experiments were also carried out accordingly. The resulting binding isotherms were analysed by nonlinear least-squares fitting of the experimental data to models corresponding to a single binding site. Analysis of the data was performed using the MicroCal Origin 7.0 software.

### NMR Spectroscopy

All NMR experiments were performed on a 600 MHz Bruker Avance III NMR spectrometer equipped with a 5 mm QCI-F cryogenic probe. NMR samples for all saturation transfer difference (STD) ^50^ experiments were prepared in 3 mm tubes filled with 170 µL of 99.9% D2O buffer containing 50 mM sodium phosphate, pH 7.4, 200 mM NaCl, 2 mM deuterated dithiothreitol (DTT), 22.2 µM 4,4-dimethyl-4-silapentane-1-sulfonic acid (DSS) and 200 µM of compound. Spectra were recorded in the presence and in the absence of 1 µM DCAF15/DDB1/DDA1 at 280 K (STD epitope mapping) and 286 K (STD Indisulam binding study). Spectra in the absence of protein were used to confirm that potential compound aggregation does not lead to false-positive STD results.

The standard Bruker pulse sequence *stddiffesgp* was used for all STD experiments. The on- and off-resonance irradiation frequencies were set to 0.33 ppm and -33 ppm, respectively. Selective saturation of the protein was achieved by a train of Sinc-shaped pulses of 50 ms length each. The total duration of the saturation periods were varied from 100 ms to 10 s (4 s for the Indisulam study). The recycling delay was set to 10 s in all experiments. The total number of scans (dummy scans) was 48 (16), a spectral width of 16 ppm was used and the number of points recorded was 32k.

^1^H-STD NMR spectra were multiplied by an exponential line-broadening function of 3 Hz prior to Fourier transformation. The on-resonance spectra were subtracted from the off-resonance spectra to obtain difference spectra, which were used for analysis.

The compound ^1^H-NMR signals were assigned by using standard small molecule NMR structure elucidation experiments (^1^H-1D, ^13^C-1D, ^1^H,^1^H-COSY, ^1^H,^13^C-HSQC, ^1^H,^13^C-HMBC). The analysis leading to the epitope map was performed using the equations described in Chatterjee et al.^50, 51^.

### Crosslinking, MALDI-MS and proteolytic digestion

The purified DDB1-DDA1-DCAF15 complex was incubated at a final concentration of 0.1 µM in 100 µL of 20 mM HEPES, pH 7.5, and 30 mM NaCl with two equivalents of RBM39(Δ150) and five equivalents of Indisulam for 2 hours at room temperature.

The crosslinking reaction was carried out with 600 equivalents of disuccinimidyl sulfoxide (DSSO, Thermo Scientific) for 1.5 hours at room temperature. This is equivalent to a lysine:DSSO molar ratio of 1:6 since DDB1-DDA1-DCAF15-RBM39(Δ150) contains 103 lysines. The covalent complex formation was analyzed by MALDI-MS prior to quenching the crosslinking reaction with NH_4_HCO_3_ to a final concentration of 20 mM (Ultraflextreme II, Bruker). The dried droplet method was used with a saturated sinapinic acid solution in CH_3_CN/H_2_O at a ratio of (75:25; v:v) with 0.1%TFA (v:v).. MALDI-MS analyses were performed in linear mode using an external calibration with the protein calibration standard II (Bruker).

The crosslinked complex was subsequently denatured with a solution of 3M urea and 180 mM NH_4_HCO_3_, and then reduced with 12 mM DTT for 1 hour at 56°C. The reduced complex was then alkylated with 36 mM iodoacetamide for 30 minutes at room temperature, and the alkylation reaction was quenched with additional 12 mM DTT. The cross-linked complex solution was diluted 3.5-fold in water and digested with trypsin (sequencing grade modified, Promega) at a 1:5 enzyme to substrate ratio (w:w) at 37°C overnight. The digestion was stopped by adding TFA at 0.1% (v:v) final concentration. The cross-linked peptides were desalted using a Sep-Pak C18 column (Waters), dried under nitrogen at 50°C, and reconstituted in HO/CH_3_CN/HCOOH (96:2:2; v:v:v) for the subsequent LC-MS^n^ analyses.

### LC-MS^n^ analysis

LC-MS^n^ data were acquired on a Lumos, Orbitrap mass spectrometer equipped with an ultra-HPLC Proxeon Easy-nLC 1200 (Thermo Scientific). Reverse phase chromatography was performed with an analytical Easy-Spray column (75µm inner diameter, 250mm length; Thermo Scientific). Cross-linked peptides were separated with a 180 min gradient from 2% to 80 % of CH_3_CN in H_2_O plus 0.1% HCOOH at a flow rate of 300nL/min. MS data were acquired with a specific DSSO-cross-linked peptide method. Briefly, MS1 was performed in the orbitrap and scanned from 300 to 1500 *m/z* with a resolution of 120,000. Only ions with charge state from 4+ to 8+ were selected for MS^2^ scans. The MS^2^ scan in the orbitrap was set to 30,000 with a precursor isolation window at 2 *m/z*. The MS^2^ normalized collision energy was fixed at 25%. MS^3^ HCD were triggered if a mass difference of 31.972 Da was observed between 2 fragment ions detected on MS2 spectrum (specific to sulfoxide MS cleavable cross-linked peptide). The two most intense ion pair ions were selected for fragmentation with a collision energy set to 30%.

### MS^n^ Data analysis and crosslink Identification

Data files were analyzed by Proteome Discoverer 2.2 (Thermo Scientific) using the XlinkX node to identify cross-linked peptides and the SEQUEST search engine for unmodified and dead-end-modified peptides. In Proteome Discoverer 2.2, the precursor mass tolerance was set to 10 ppm, the MS^2^ filter for peptide tolerance at 20 ppm, and the MS^3^ peptide fragment tolerance at 0.6 Da. Data was searched with a 1% FDR criteria against a restricted database containing the 4 proteins (DCAF15, DDB1, DDA1, and RBM39). Crosslinked peptides identified with Proteome Discoverer were filtered for a confident identification with Xlink score greater than 50. The protein-protein interaction mapping for the complex was visualized with XiNET Viewer^52^.

### Crystallization and structure solution of the DCAF15-DDB1(ΔBPB)-DDA1-RBM39(RRM2)-Indisulam complex

Initial crystallization screens comprised of 1800 different crystallization conditions utilizing DCAF15-DDB1-DDA1-RBM39(RRM2)-ligand complexes identified crystals with excellent hexagonal morphology. However, these hexagonal crystals were extremely soft and exhibited poor diffraction (approximately 6-8 Å) and extreme radiation sensitivity. These crystals could not be optimized to yield higher resolution diffraction. Initial molecular replacement solutions using a DDB1 search model (PDB accession code 3E0C) and the observation of radiation sensitivity strongly suggested that the crystals had a high solvent content due to the packing arrangement enforced by full-length DDB1 and that the B-propeller domain (BPB) was positioned at a different angle in the complex versus the apo-structure. Also, the initial EM maps showed that the B-propeller domain showed increased flexibility versus the structure of apo-DDB1. Given these observations, a construct in which the BPB domain was deleted was utilized to yield an approximately spherical complex with more possible packing arrangements, as well as reduced flexibility. This protein engineering yielded a well-behaved protein complex that forms crystals with improved robustness and diffraction; these crystals typically diffract to 2.3-2.8 Å.

DCAF15-DDB1(ΔBPB)-DDA1-RBM39(RRM2)-ligand complexes were crystallized as follows. DCAF15-DDB1(ΔBPB)-DDA1 at a concentration of 10.0 mg/ml was combined with 1.8 molar equivalents of both RBM39(RRM2) and ligand, respectively. This mixture was incubated on ice for 30 minutes to allow the complex to form and was then spun in an ultracentrifuge at 14,000 rpm for 10 minutes to remove debris and aggregates. Crystallization trays were set up using the hanging drop method in INTELLI-plates using 0.2 ul of protein solution and 0.2 ul of precipitant. A precipitant grid screen consisting of 2% (v:v) Tacsimate^TM^, pH 5.0, 0.1 M sodium citrate tribasic dihydrate, pH 5.6, and 10-20% (w:v) polyethylene glycol 3350, was used. Crystals grew at 18°C in 5 days. Crystals were cryo-protected using well solution supplemented with 25% v:v of glycerol. All data was collected at the Advanced Light Source on beamline 5.0.2 using standard collection protocols at a wavelength of 1 Å, which provided the highest flux. Several distinct and disparate areas of single crystals were used for data collection, and these arcs were combined to yield data sets of high multiplicity.

The DCAF15-DDB1(ΔBPB)-DDA1-RBM39(RRM2)-Indisulam co-structure was solved using a hybrid molecular replacement/pseudo-atom approach. An initial molecular replacement solution was executed using an appropriately-pruned search model consisting of the crystal structure of apo-DDB1 (PDB accession code 3E0C) and the NMR structure of RBM39 RRM2 (PDB accession code 2JRS), in the given order. This molecular replacement solution was carefully refined against the X-ray data using a resolution cut-off of all data with an I/σ greater than or equal to 1, corresponding to a resolution cut-off of 2.50 Å, using cross-validation via Rfree to monitor the suitability of the refinement. Initial electron density maps showed new features and improved information content for both DDB1 and RBM39 (such as different sidechain positions) which were not present in the refined search models. These structures were refined to convergence. At this point, 32% of the mass of the complex was present in the model and the electron density maps showed some features which indicated that DCAF15 was bound. It was possible to visualize the docking helix density and there was noisy density was present for a few β-strands, but these could not be accurately placed, nor could the maps be improved with other standard methods alone, such as solvent-flattening. To prevent having to produce selenomethione-labeled protein, a pseudo-atom approach was used to provide additional phase information. Using the Phenix suite^53^, the current electron density maps were computationally interrogated in the presence of the already refined DDB1-ΔBPB and RBM39-RRM2 structures and at each position where the electron density map showed a peak at 1σ in the 2Fo-Fc map and at 2σ in the Fo-Fc map, a water molecule was placed. The role of these water molecules was to provide a scattering surrogate for other atoms in both main-chain and side-chains. To allow these pseudo-atoms to more effectively mimic atomic centers, the effective VDW radii was decreased and the real-space correlation cutoff used in the atom placement was effectively disabled. This DDB1-ΔBPB:RBM39-RRM2:pseudo-atom model was carefully refined to prevent over-fitting (again via observation of Rfree), followed by additional placement of pseudo-atoms and refinement of the model until convergence had been reached. To minimize bias, the electron-density maps from this step were subjected to both solvent-flattening and histogram matching, and the positions of all pseudo-atoms were visually inspected against these electron density maps and appropriately repositioned and/or deleted. This adjusted model, which consisted of a DDB1-ΔBPB:RBM39-RRM2:pseudo-atom ‘complex’, had an R/Rfree of approximately 28%/35% and showed numerous additional features, including the majority of the β-strands which comprise the body of DCAF15, as well as the chain of DDA1. The electron-density map from this model was again subjected to solvent-flattening and histogram matching and used by SOLVE to computationally build a skeleton for the protein as well as assignment of amino acid sequence where possible. These maps also unambiguously identified the binding site of Indisulam as well as all bordering amino acids, which were fit. The skeleton coordinates, as well as order/disorder and secondary structure predictions were then used to build the remainder of both DCAF15 and DDA1 using standard 2Fo-Fc and Fo-Fc maps, as well as ‘feature-enhanced’ maps. The structure of the complex was consistent with the cross-linking data and EM maps. The structure of the complex was then refined to convergence via multiple cycles of manual rebuilding and refinement using data from 69.10 -2.3 Angstroms (consistent with a CC1/2 cutoff of 0.493 for the high-resolution data) using both the Phenix^53^ and BUSTER^54^ program suites. The final structure of the complex had an R of 20.4% and an Rfree of 24.9%. The final crystal co-structure consists of 1,392 residues, Indisulam, a glycerol molecule, and 875 waters. The co-structure has a clashscore of 10.65. The model has 93.78% of the protein residues in the favored region of the Ramachandran plot, 6.00% in the allowed region, and 0.22% as outliers. The Molprobity score is 2.28^55^; there are extremely regions at the periphery of DCAF15 where it is difficult to fit appropriate rotamers.

Please note that to execute such a phasing strategy, it is necessary to have complete, well-measured data, particularly for a complex of this size. The DCAF15-DDB1(ΔBPB)-DDA1-RBM39(RRM2)-Indisulam co-structure was subsequently used to solve the DCAF15-DDB1(ΔBPB)-DDA1-RBM39(RRM2) co-structure with compound **7** by molecular replacement.

### Cryo-EM sample preparation and data acquisition

Two µM of the DCAF15-DDB1-DDA1 complex was incubated with 200 µM of Indisulam and 10 µM of RBM39(RRM2) at 4°C for 30 minutes. 4 μL aliquots of the complex were applied to glow-discharged, 300-mesh Quantifoil R 1.2/1.3 grids (Quantifoil, Micro Tools GmbH). These grids were blotted for 3 seconds and subsequently plunged into liquid ethane using an FEI Mark IV Vitrobot operated at 4°C and 90% humidity. High-resolution images were collected with a Cs-corrected FEI Titan Krios TEM operated at 300 kV equipped with a Quantum-LS Gatan Image Filter and recorded on a K2-Summit direct electron detector (Gatan GmbH). Images were acquired automatically (with EPU, ThermoFisher) in an electron-counting mode, using a calibrated magnification of 58140 corresponding to a magnified pixel size of 0.86 Å. Exposures of 12 seconds were dose-fractionated into 40 frames. The total exposure dose was ∼50 e-/Å^2^. Defocus values per frame varied from -0.8 to -2.4 μm.

### Image processing of cryo-EM data

The collected frames were processed using cisTEM^56^. Whole-frame motions were corrected, followed by estimation of the contrast transfer function (CTF) parameters. Images with CTF fits to 4 Å or better were selected. 450,000 coordinates were then automatically selected based on an empirical evaluation of maximum particle radius (70 Å), characteristic particle radius (50 Å), and threshold peak height (4 standard deviations above noise). Three rounds of 2D classification into 50 classes were performed to remove false positives and suboptimal particles. The remaining 150,371 particles were used for ab initio 3D reconstruction with applied C1 symmetry. These particles were subsequently used for iterative 3D classification and auto refinement, which resulted in an approximately 3.3 Å resolution map. Maps were auto-sharpened by using the Phenix suite^53^. The resolution values reported are based on the gold-standard Fourier shell correlation curve (FSC) criterion of resolution cutoff of 0.143. Simultaneously, a total of 415203 particles were extracted for processing using the Relion 3 software package^57^. Particle sorting included two cycles of reference-free 2D classification. The 386408 particles in the best 2D classes were used for 3D refinement. The generation of initial model was carried out in cisTEM^56^. 3D classification was performed without alignment to separate different conformational states (Relion3). Auto-refinement of particles with a soft mask (relion_create_mask) around complex resulted in a 3.54 Å resolution map. The crystal structure of DDB1 (PDB accession code 5JK7) was manually fitted into the cryo-EM map using Coot^58^. The DDA1 density was located using in-house cross-linking data and DDA1 model was built with a partially assigned sequence. The NMR structure of the RBM39 RRM2 domain (PDB accession code 2JRS) was manually fitted to the cryo-EM map. Indisulam density could be located at the interface of DCAF15 and RBM39 domain. To complete the model of DCAF15, initially secondary structures were placed into the EM maps using poly-alanine α-helices and β-sheets. Sequence assignments of the observed secondary structure were then completed using the crystal structure of the DCAF15 complex and side-chains were added as appropriate. This atomic model was subjected to multiple cycles of model rebuilding using Coot and real space refinement against the map using Phenix^53^. This process resulted in an atomic model of the ternary complex that fits well into the cryo-EM density map. The final EM co-structure consists of 1,308 residues and Indisulam. The co-structure has a clashscore of 9.00^55^. The model has 88.77% of the protein residues in the favored region of the Ramachandran plot, 10.21% in the allowed region, and 1.03% as outliers.

### Differential scanning fluorimetry

Thermal shift assays were performed with 5 uM of purified DCAF15-DDB1-DDA1 or DCAF15-DDB1 complex in buffer D (see purification protocol). The samples were mixed with 5x SYPRO Orange (Molecular Probes) prior to the thermal cycle. The temperature was ramped from 4 to 95 °C in a ViiA7 real-time PCR machine (Applied Biosystems). The protein melting temperature, Tm, was calculated based on the resulting fluorescence data using curve-fitting to a Boltzmann function by the Protein Thermal Shift software version 1.1 (Life Technologies). The standard deviation was calculated by comparing three replicate experiments. Data were plotted using GraphPad Prism 8.

### Cell culture

HCT116 and HEK293T cells were obtained by ATCC. HCT-116 cells were maintained in McCoy’s 5A and HEK293T were maintained in DMEM (Invitrogen), both media supplemented with 10% (v:v) FBS, 1% (w:v) penicillin-streptomycin, and 2 mM L-glutamine (VWR) in a humidified incubator held at 37⁰C and 5% CO2. Cells were confirmed to be mycoplasma negative via MycoAlert^TM^ Mycoplasma Detection Kit (Lonza).

### Cell viability assays

HCT116 or HEK293T cells were trypsinized, diluted in growth media to a final concentration 2.5x10^4^ cells/mL, and plated in 384-well plates (Corning #3707) at 40 µL/well. Cells were incubated overnight at 37 °C (5% CO2). Test compounds were diluted to various concentrations in DMSO and further diluted 40X in growth media. Cells were treated with 10 µL/well of diluted of test compound or vehicle (0.05% DMSO final concentration) via BioMeK liquid handled and incubated for 72hrs at 37C (5%CO2).

After 72 hr incubation, treated cells were equilibrated to room temperature for 30 min. <u>CellTiter-Glo®</u> reagent (Promega) was added at 20 µL per well and plates were placed on an orbital shaker for 30 s prior to a 10 min incubation at room temperature. Luminescence was measured using an Envision MultiLabel reader (200 ms read time). Readings from all DMSO wells were averaged and each test well reading (compound treated) was normalized to DMSO. Results were plotted in GraphPad Prism 8 and data were fit using the nonlinear fit module (“3 parameter – log dose vs response”) to determine IC50.

### siRNA knockdown of DDA-1

For siRNA transfection, 10^6^ cells plated per well of a 6-well plate were transfected with 150 pmol siRNAs (Ambion: Negative Control #AM4611, DDA1-2 Cat #4392420 ID #s35423) using 9 uL of Lipofectamine RNAiMAX (Life Technologies). After 40 hours post-transfection, cells were washed with PBS and re-suspended in DMEM, and then treated with DMSO or Indisulam for another 24 hours before cell collection. Cells were pelleted and washed with 2 x 1mL PBS and frozen at -80°C overnight and then thawed and lysed with 100 µL of RIPA buffer (Thermo Scientific) supplemented with 1X HALT protease inhibitor cocktail (Thermo Scientific).

### Western blotting and antibodies

LDS samples were prepared using LDS sample buffer (Invitrogen, REF #NP0007) and Sample Reducing Agent (Invitrogen, Cat #NP0009) and separated using Bio-Rad PowerPac HC system using NUPAGE 4-12% Bis-Tris gels (Invitrogen, Cat #NP0323BOX). Proteins were transferred via Bio-Rad Trans-Blot Turbo transfer system onto Bio-Rad Trans-Blot turbo transfer pack with 0.2 um nitrocellulose membranes (Invitrogen, Cat #1704158).

Blots were incubated with primary antibody solutions made in TBS-T with 5% milk for RBM39 (Sigma, Cat #HPA001591, 1:2500) GAPDH (CST, Cat #2118L, 1:1000), Vinculin (Cell Signaling Technology, Cat# 13901S, 1:1000), ZNF277 (Pro-Sci, Cat #46-616, 1:1000), RBM23 (Invitrogen, Cat# PA5-52060, 1:1000), or DDA1 (Proteintech, Cat #14995-1-AP, 1:1000) overnight at 4°C. Blots were then washed 3 x 5 mL TBS-T and incubated with secondary antibody (EMD Millipore, Cat #AP307P, 1:2500) for 1h at 25°C, then visualized with Amersham ECL (GE Life Sciences, Cat #RPN2236) and imaged using Amersham Hyperfilm ECL (GE Life Sciences, Cat #28906839).

### Fluorescence polarization assays

To study the impact of various RBM39 substitutions on aryl sulfonamide-induced recruitment to DCAF15, we leveraged the differential binding affinity of a FITC-labeled Indisulam analog to DCAF15 complex-alone versus a DCAF15-RBM39 complex in a fluorescence polarization assay.

We first determined the concentration of an Avi-DCAF15-DDB1-DDA1 complex required to yield probe binding (high polarization) in the presence of RBM39(Δ150), signaling ternary complex formation, and low probe binding (low polarization) in the absence of RBM39(Δ150). Avi-DCAF15/DDB1/DDA1 was diluted to 20 µM in FP Buffer, consisting of 20 mM HEPES pH 7.2, 150 mM NaCl, and 0.01% (v:v) Tween-20, and serially diluted in the presence or absence of 2 uM His-ZZ-RBM39(Δ150). Each mixture was dispensed into a black 384-well plate (Corning #3575) at 10 µM per well. Compound **9** (FITC-probe) was diluted to 40 nM in FP Buffer and dispensed at 10 µL per well to yield a final concentration of 20 nM. Plates were incubated at room temperature for 1 hr and read on an Envision MultiLabel reader equipped with standard FITC-FP protocol/mirror sets. The Envision Assay Optimization Wizard was used to adjust detector gain and to determine G-factor (measured on compound **9**-only wells) for mP calculation.

To assess the impact of RBM39 mutation on ternary complex formation with DCAF15 and compound **9**, we titrated RBM39 variants in the presence of compound **9** and DCAF15, measuring fluorescence polarization. Avi-DCAF15/DDB1/DDA1 and compound **9** were diluted to 200 nM and 40 nM in FP buffer, respectively, and dispensed into black 384-well plates (Corning #3575) at 10 µL per well. RBM39 variants were diluted in FP buffer to 20 µM each and serially diluted, with each mixture then added at 10 µL per well to yield final volume of 20 µL. Plates were incubated at room temperature for 1 hr and read on an Envision MultiLabel reader as before.

### TR-FRET recruitment assays

In TR-FRET buffer consisting of 20 mM HEPES, pH 7.4, 150 mM NaCl, and 0.05% (v:v) Triton X-100, a solution of 20 nM LanthaScreen™ Tb-Streptavidin (Thermo Scientific, Cat# PV3965), 150 nM biotinylated-Avi-DCAF15/DDB1/DDA1, 500 nM 6XHis-ZZ-RBM39(Δ150), and 50 nM anti-6xHis-FITC (AbCam, Cat# ab1206) was prepared and transferred into a black, 384-well Corning plate (#3575) at 20 uL per well. DMSO stock solutions of various compounds and respective dilutions were transferred acoustically via Echo 555 Liquid Handler (Labcyte, Inc) at 100 nL per well. After transfer, plates were incubated at room temperature for 1 hr and TR-FRET was read on the Envision Multi-label reader (using the following conditions).

|  |  |
| --- | --- |
| Light source | Flash lamp |
| Top mirror | LANCE/DELFIA |
| Bottom mirror | N/A |
| Exc. filter | UV2 (TRF) 320 |
| Using of excitation filter | Top |
| Ems. filter | Emission 520 |
| Using of emission filter | Top |
|  | Photometric |
| 2nd ems. filter | 492 |
| Using of 2nd emission filter | Top |
| Measurement height | 6.5 mm |
| Cycle | 2000 |
| Delay | 60 |
| Number of flashes | 100 |
| Number of flashes for 2nd detector | 100 |
| Number of sequence windows | 1 |
| Total time of windows | 300 |
| Excitation light | 100% |

TR-FRET ratios (520 nm /490 nm emission signals) were analyzed using Excel (calculating averages and standard deviations) and these data were plotted in GraphPad Prism 8.

### Proteome-wide motif search

Given the complementarity between the RBM39(T262-P272) alpha helix, Indisulam, and DCAF15 in the ternary complex, we queried the proteome for similar alpha helicies, bearing residues critical for DCAF15-Indisulam binding, to identify additional proteins that may be recruited to DCAF15. From described RBM39 mutagenesis studies and previously published work^17, 18^, we determined residues M265, G268, E271, and P272 to be most critical to DCAF15-Indisulam binding and established the “X^1^XXM^4^XXG^7^XXEP^11^” motif as a putative degron. To identify compatible proteins with this motif, we first accessed 20,407 unique human protein sequences from Uniprot (www.uniprot.org), cataloging the availability of any associated x-ray/NMR structures (6,475 entries). For proteins with available structures, we analyzed for the presence of an alpha helix with a glycine residue and aligned this region on the alpha helix of RBM39 in the ternary structure model with DCAF15. A backbone alpha carbon RMSD of each protein structure versus the RBM39(RRM2) structure was calculated and if the RMSD was <2Å (as in 3112 structures), we surveyed for steric clashes with DCAF15. A steric clash of less than 10 heavy atoms between DCAF15 and the target protein was considered acceptable. Proteins with low RMSD (<2Å) and minimal steric clash to DCAF15 were considered. Finally, hit sequences were filtered by the presence of the “X^1^XXM^4^XXG^7^XXEP^11^” sequence.

### Expression proteomics

TMT-based expression proteomics was performed as previously described^59^ with few modifications. Indisulam-treated HCT116 cells (10^6^ cells treated with 10 uM Indisulam for 4 hrs.) were harvested, washed 3 times with PBS, lysed with 500 µL lysis buffer (8M Urea, 1% SDS, 50 mM Tris pH 8.5, with protease and phosphatase inhibitors added), and then sonicated to shear DNA aggregates. After centrifugation, protein concentrations were measured by Micro BCA™ Protein Assay Kit (Thermo Cat#23235).

For each sample, 200 µg protein was aliquoted and reduced with 5 mM Dithiothreitol (DTT) for 1 hr at room temperature (RT), alkylated with 15 mM iodoacetamide (IAA) for 1 hr at RT in the dark, and then quenched with 10 mM DTT for 15 min at RT. Alkylated proteins were purified via chloroform/methanol precipitation^60^, dissolved in denaturing buffer (8 M urea, 50 mM Tris, pH 8.5), and diluted with seven volumes of 50 mM Tris, pH 8.5. Protein was digested using Trypsin/Lys-C mix in an enzyme to protein ratio of 1:25 and incubated overnight at 37 °C. A second digestion was performed with additional Trypsin/Lys-C mix (enzyme to protein ratio of 1:50) for 4 hours.

The peptide sample was then desalted using a Water’s tC18 SepPak plate (Waters Cat# 186002321), dried down, and then resuspended in 100 µL 0.1 M TEAB buffer, pH 8.5. Peptide concentrations were determined using the Pierce™ Quantitative Fluorometric Peptide Assay (Thermo Cat#23290) and normalized between samples (∼2 mg/mL)

For each sample, 200 ug peptides were labeled via TMT10plex™ Isobaric Label Reagent kit (Thermo Cat#90111) at the ratio of 4 units of TMT reagent to 1 unit of peptide. TMT labeling efficiency was checked by running an MS analysis. Once the labeling efficiency was confirmed to be greater than 99%, the reaction was quenched with 0.5% with hydroxylamine for 15 min at RT. Equal amounts of each TMT-labeled sample were combined, desalted using Water’s tC18 SepPak plate (Waters Cat#186002321), and then fractionated by HPLC - Waters XBridge C18 (3.5 µm, 300 x 4.6 mm) column with gradient of 10-40% mobile phase B (90% acetonitrile with 5 mM ammonium formate, pH 10) in mobile phase A (5 mM ammonium formate, 2% acetonitrile). Final fractionated peptide material was pooled into 24 fractions (∼1-2 µg peptides/fraction).

Each fraction was analyzed using an Orbitrap Fusion™ Lumos™ Mass Spectrometer (Thermo) equipped with a Reprosil-PUR column (1.9 µm beads, 75 µm ID x 15 µm tip x 20 cm, 120 Å). Samples were run using gradients of 7-28% mobile phase B (80% acetonitrile with 0.1% formic acid) in mobile phase A (0.1% formic acid) using the SPS MS3 mode. Thermo Proteome Discoverer™ was used to analyze raw data and determine major cutoff parameters for peptide quantification (i.e. precursor contamination <50%, minimum average reporter ion with signal/noise >10, and peptide-spectrum match (PSM) ≥1 for all peptides). Custom iPython notebook processing with limma statistical analysis and normalization was used to determine fold-changes and p-values between duplicate DMSO- and Indisulam-treated samples. Specifically, the scipy.stats.f_oneway function was used to perform one-way ANOVA and generate F-statistics and associate p-values from the F-distribution. Samples were assumed to be independent, normally distributed, and homoscedastic.

### Chemical synthesis and characterization

All chemical synthesis procedures and characterization data are provided in Supplementary Note 2.

### Statistical Analysis

All statistical analyses were performed using Prism 8 (GraphPad) unless otherwise stated in the Methods section.

## Acknowledgments

The authors thank M. Renatus for providing DCAF15 constructs and M. Li for providing DDB1 and DDB1(ΔBPB) constructs. We also thank G. Pardee for baculovirus generation and protein expression, S. Widger for additional expression support, and X. Ma for helpful discussions on ligase structural biology. Finally, we thank J. Bradner, J. Shulok, R. Jain, J. Porter, and J. Tallarico for helpful discussions and input on this manuscript.

## Author contributions

R.B., A.F., J.Z., B.O., S.J., and P.M. designed and/or synthesized reported compounds. N.C., P.G., and H.V. performed crosslinking/mass-spectrometry studies. A.B., D.K., A.G, performed surface plasmon resonance experiments, while D.K. performed analytical ultracentrifugation. V.H. and R.G.K. performed proteome-wide motif searches and structural/computational modelling. C.B., H.S., and C.W. collected and processed Cryo-EM data. F.X. and J.C. conducted expression proteomics experiments. A.O.F. and A.F. performed biological NMR experiments. W.S. performed crystallographic screening and crystal optimization; D.E.B. designed protein constructs, collected X-ray crystallography datasets, reduced data, determined initial crystal structures and refined final structures; M.K. refined final structures. L.X. purified DCAF15 complexes and RBM39 variants, and performed ITC and DSF experiments. J.P. and A.B. purified RBM39 variants, performed FP and TR-FRET assays, cellular viability assays, siRNA knockdown, and western blots. All authors contributed to writing. D.E.B., J.S., and J.P, wrote and edited the final manuscript. D.E.B., J.S., L.X., and J.P contributed intellectual and strategic input.

## Competing interests

All authors are employees of Novartis, or were at the time of this study.

## Data availability

The authors declare that the data supporting the findings of this study are available within the publication and its Supplementary Information files or have been deposited in the RCSB Protein Data Bank (PDB, http://www.rcsb.org) or Electron Microscopy Data Bank (EMDB, http://www.ebi.ac.uk/pdbe/emdb/), as appropriate. The PDB accession code for the human DCAF15-DDB1-DDA1-RBM39(RRM2)-Indisulam co-structure is ----. The EMDB accession code for the human DCAF15-DDB1-DDA1-RBM39(RRM2)-Indisulam co-structure is ----.

---

**Supplementary Figure 1.**
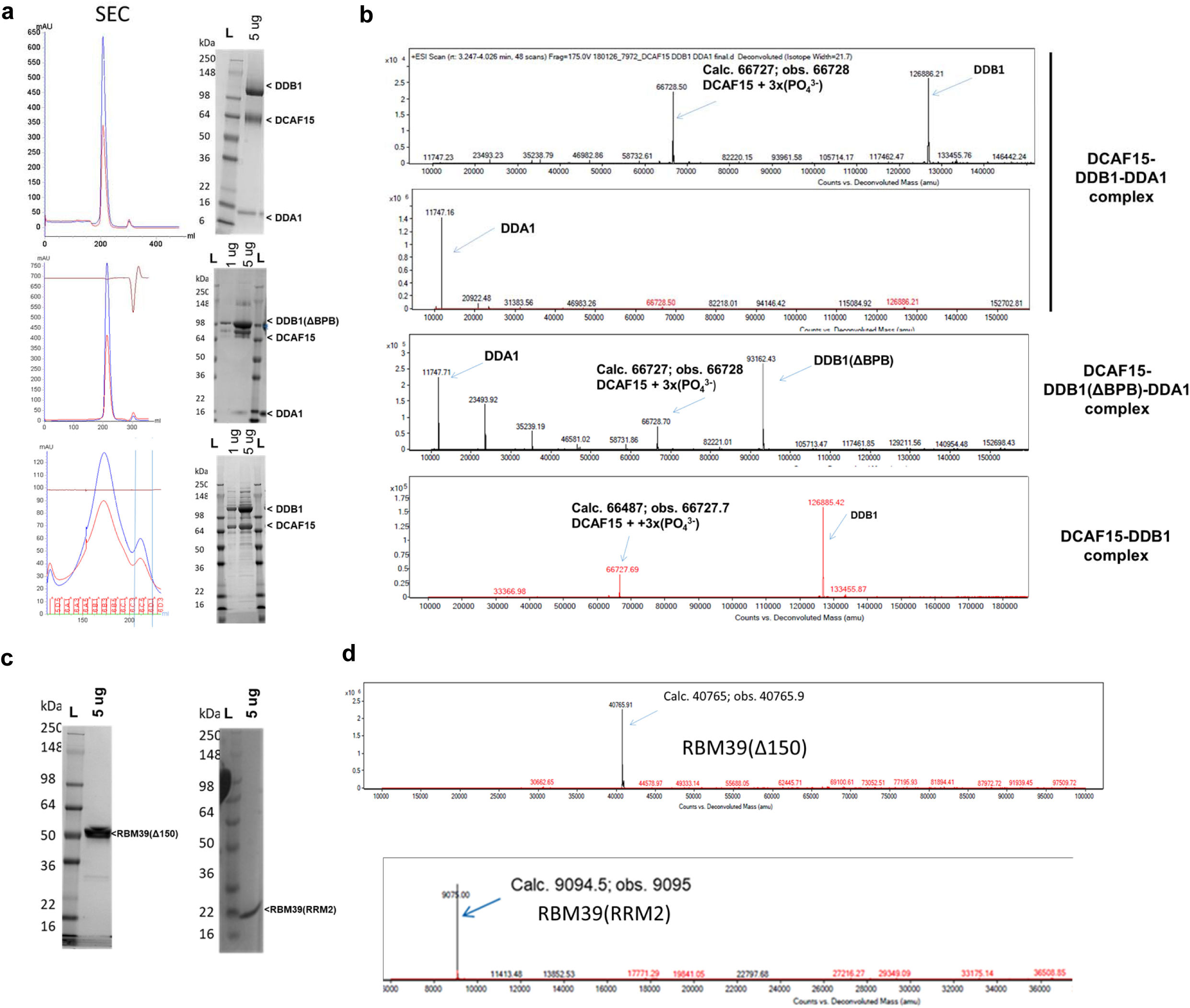
Characterization of purified RBM39 and DCAF15-DDB1-DDA1 complexes. **a**, Size-exclusion chromatography (SEC) traces (absorbance at 280 nM) from GE Superdex 200 column separation of purified DCAF15-DDB1-DDA1 (top), DCAF15-DDB1(ΔBPB)-DDA1(middle), and DCAF15-DDB1 (bottom) samples alongside Coomassie-stained SDS-PAGE analysis. **b**, LC-MS analysis and mass determination for purified DCAF15- DDB1-DDA1 (top), DCAF15-DDB1(ΔBPB)-DDA1(middle), and DCAF15-DDB1 (bottom) samples. **c**, Coomassie-stained gels for RBM39(Δ150) and RBM39(RRM2) proteins (left and right, respectively) and **d**, associated LC-MS analysis to confirm masses (top and bottom, respectively)

**Supplementary Figure 2.**
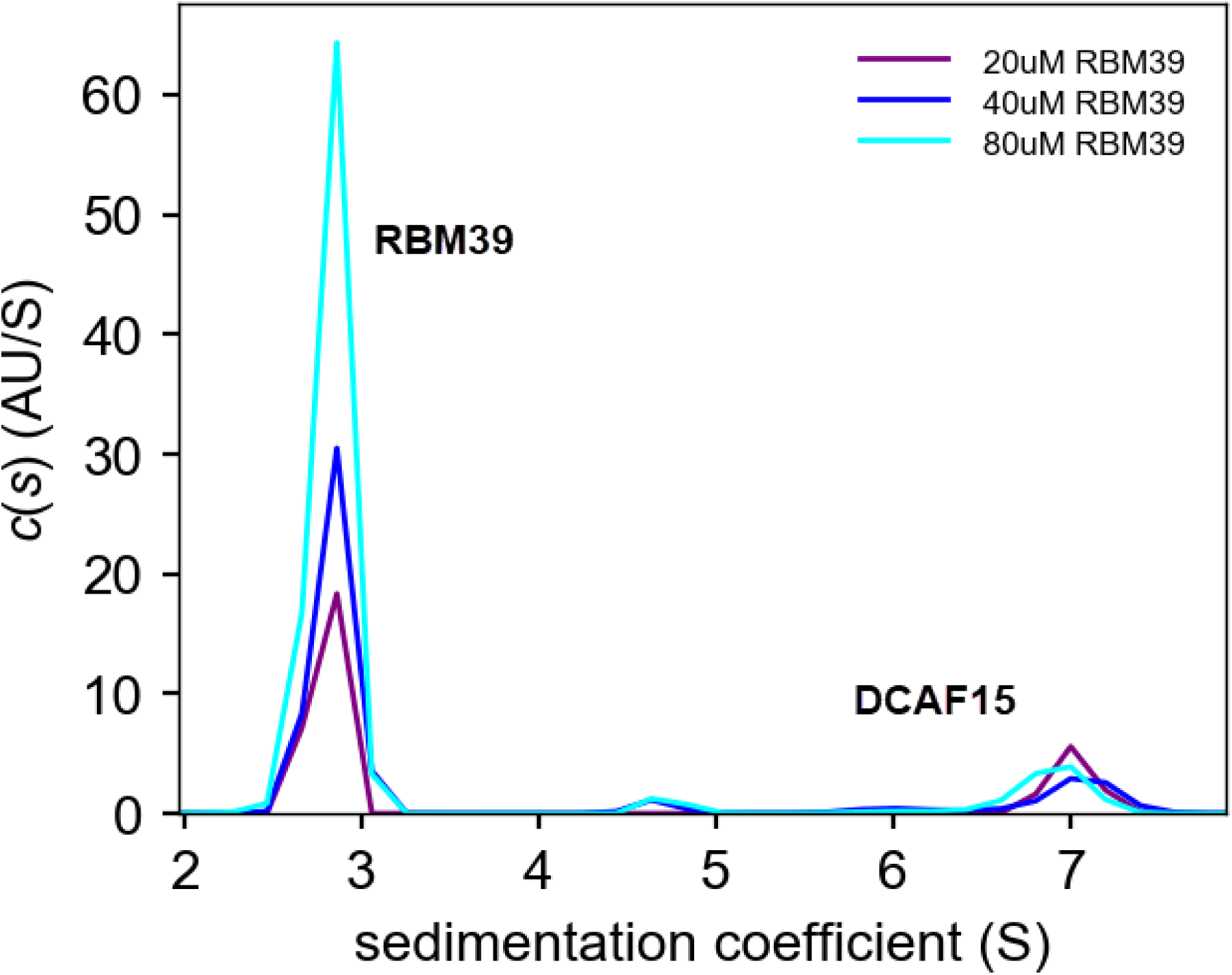
Recruitment of RBM39 to DCAF15 is Indisulam-Dependent. 2.5 µM DCAF15-DDB1(ΔBPB)-DDA1 (DCAF15) was incubated with increasing concentrations of His-ZZ-RBM39(Δ150) (RBM39) as listed in the legend. The components were subjected to sedimentation velocity in 50 mM HEPES (7.5), 300 mM NaCl & 1 mM TCEP @ 42,000 rpm for 5 h, 20 C. The sedimentation coefficient distribution displays independently migrating His-ZZ- RBM39(Δ150) (2.7 S) & DCAF15-DDB1(ΔBPB)-DDA1 (6.8 S). In the absence of Indisulam, concentrations as high as 80 μM His-ZZ-RBM39(Δ150) do not display a dose-dependent increase the integrated area of a putative DCAF15-DDB1(ΔBPB)-DDA1-RBM39 complex peak or migrate with a higher sedimentation coefficient, inconsistent with the formation of a stable complex in the absence of Indisulam.

**Supplementary Figure 3.**
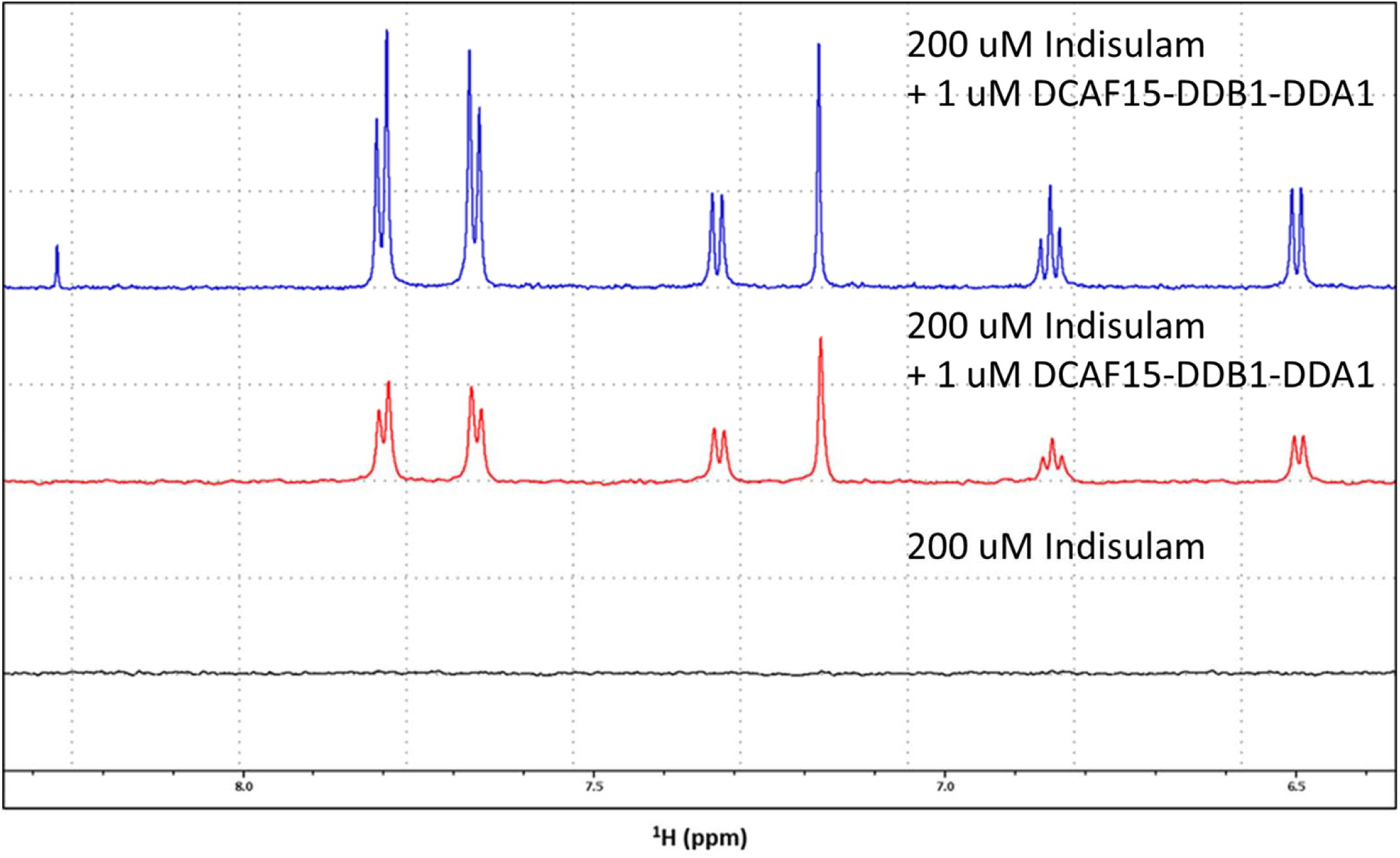
^1^H-STD Indisulam spectra in the presence and absence of DCAF15-DDB1-DDA1. Evidence of binding of Indisulam to DCAF15/DDB1/DDA1: ^1^H-1D spectrum of 200 µM Indisulam in the presence of 1 µM DCAF15/DDB1/DDA1 (blue); replicate ^1^H-STD spectrum of the same solution (red); ^1^H-STD spectrum of a solution containing 200 µM Indisulam, but no protein (black).

**Supplementary Figure 4.**
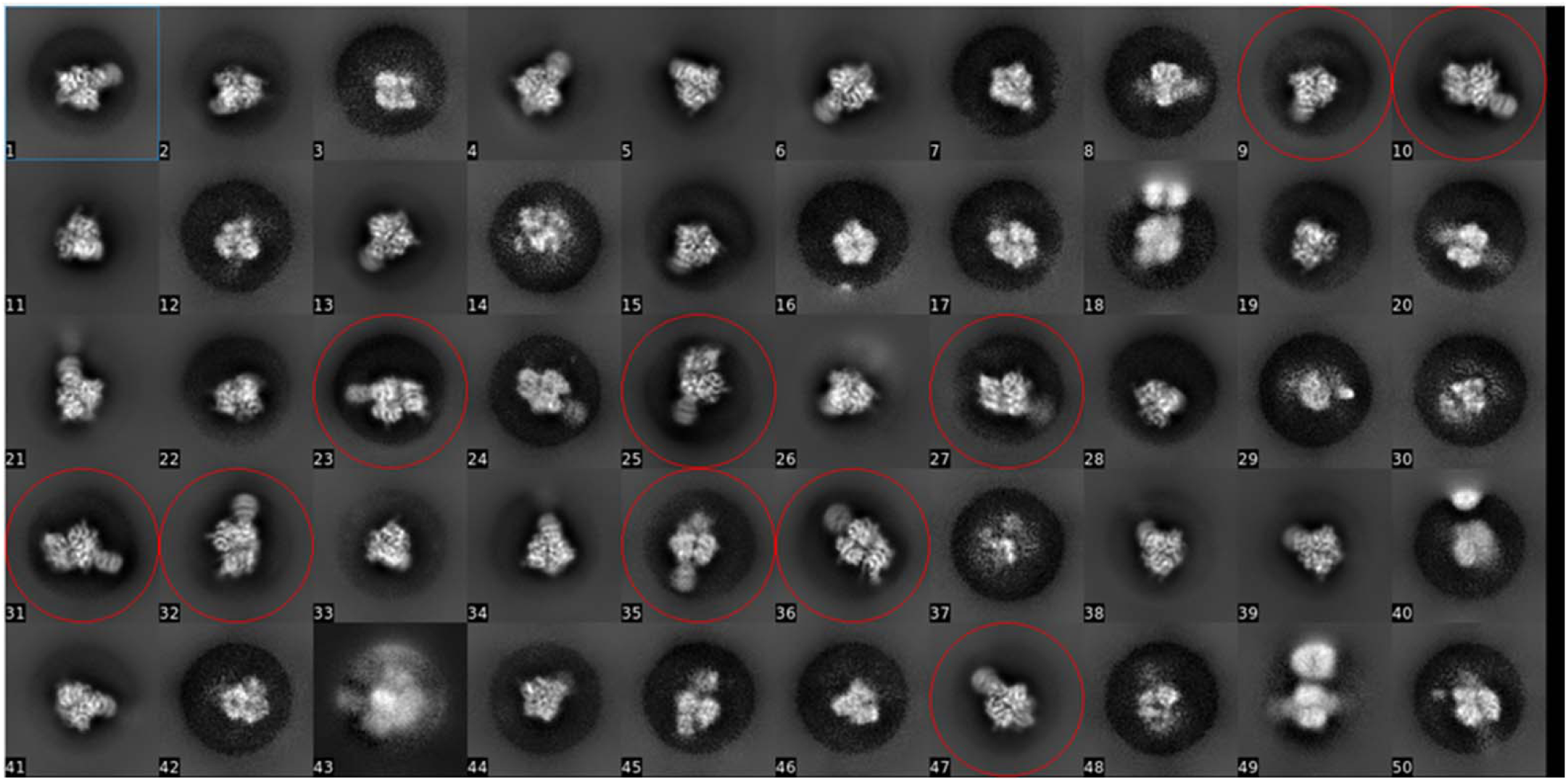
EM 2D-class averages show mixture of DCAF15 ternary complexes and DDB1 alone. 2D-class averages showing a mixture of ternary complex and DDB1 alone. For 3D-classification, only ternary complex class, marked with red circle were chosen for further processing.

**Supplementary Figure 5.**
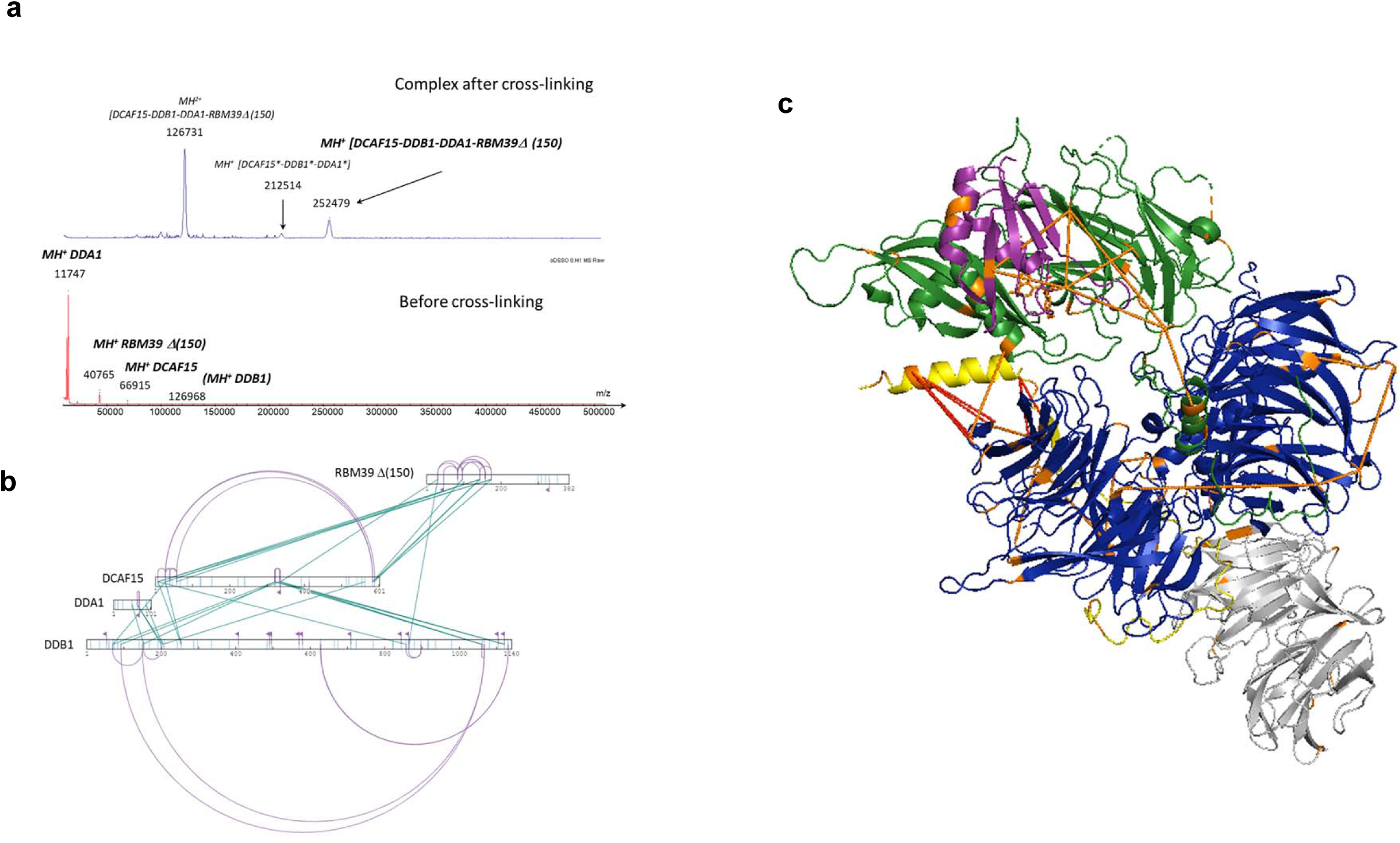
MS crosslinking studies to map DCAF15-DDB1-DDA1-Indisulam-RBM39(Δ150) interactions. **a**, MALDI MS spectra of the complex before (bottom) and after (top) the DSSO crosslinking reaction.. The major species at 252 K_d_a confirms the covalent crosslinking of the four protein components with a stoichiometry 1:1:1:1 for DCAF15-DDB1-DDA1-Indisulam-RBM39(Δ150). **b**, XiNET visualization of the protein-protein interaction mapping within DCAF15-DDB1-DDA1-Indisulam-RBM39(Δ150): 30 inter-protein (green) and 30 intra- (purple) protein crosslinks were confidently identified. **c**, PyMOL visualization of the cryo-EM structure with the identified inter-protein crosslinks (orange) in the DCAF15-DDB1-DDA1-Indisulam-RBM39(Δ150) complex. Inter-protein crosslinks in red enabled the determination of DDA1 positioning at the interfaces of DCAF15 and DDB1. Six protein crosslinks were identified between the RBM39 (RRM2) domain and DCAF15. Five protein crosslinks were identified between DDB1 and DDA1 and three of these provided important spatial constraints for the EM model. These three protein crosslinks were between DDB1(Lys153) and DDA1(Lys51), DDB1(Lys200) and DDA1(Lys65), and DDB1(Lys204) and DDA1(Lys65). Two protein crosslinks were identified between the unresolved C-terminal (69-102) of DDA1 and DDB1. Twelve protein crosslinks were also identified to disordered areas of DCAF15, showing interactions between DCAF15 and the N- and C-terminal of DDB1.

**Supplementary Figure 6.**
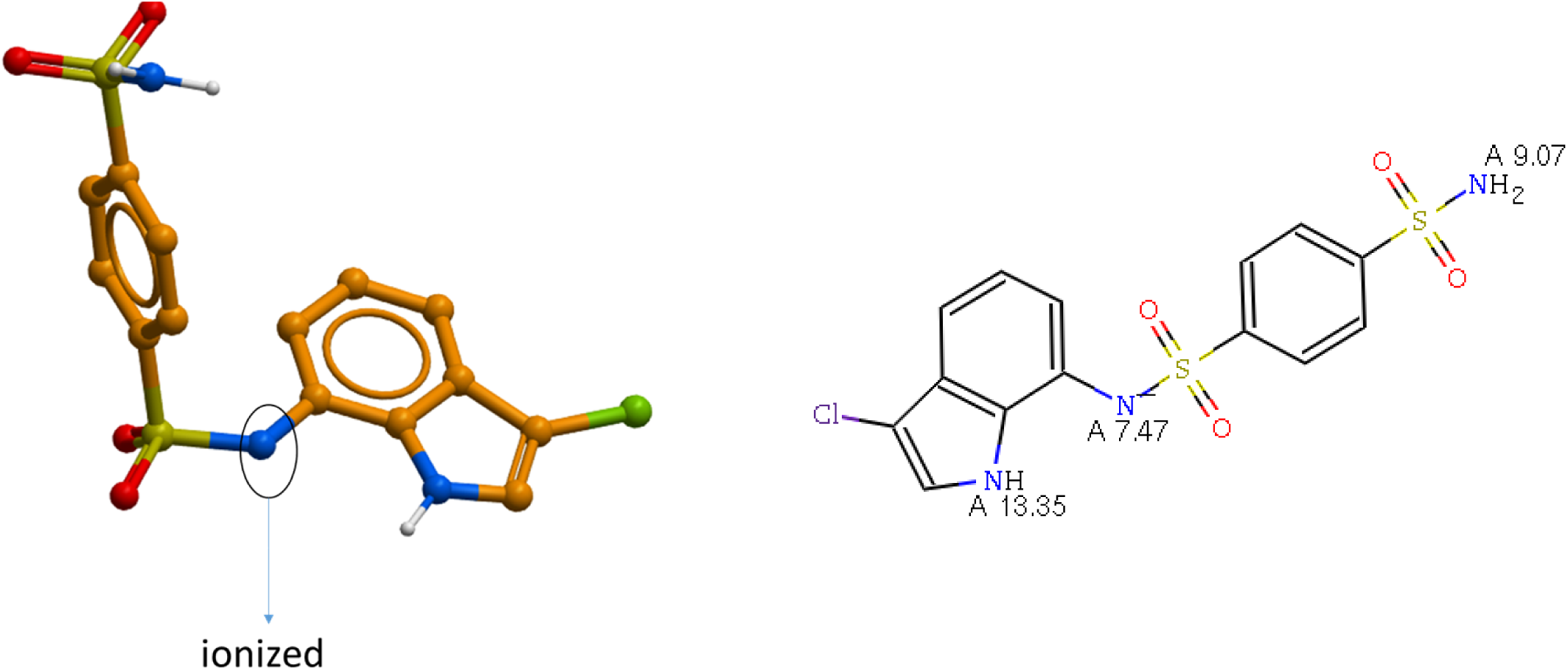
Indisulam’s DCAF15-bound geometry represents a low energy conformer assuming an ionized state. Predicted pKa value of the sulfonamide nitrogen of Indisulam is 7.47 (MoKa program), suggesting a 50% probability for compound ionization at pH of 7.4. The DCAF15-RBM39-bound geometry of the ionized species predicted by the X-ray structure is a favored low energy conformer.

**Supplementary Figure 7.**
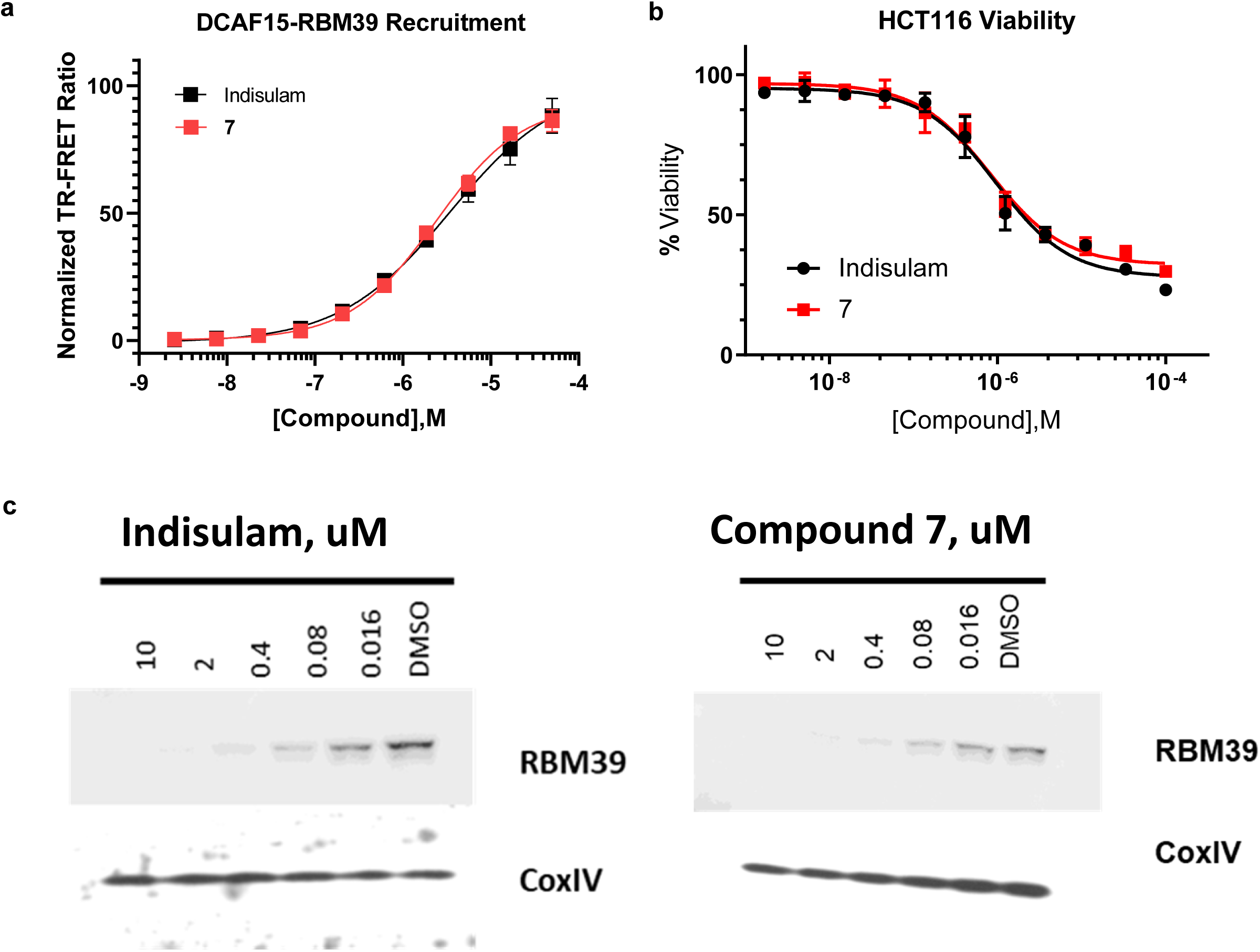
Biochemical and cellular activity of compound 7 on DCAF15-dependent RBM39 recruitment and degradation. **a**, Select data from Fig. 5 reproduced for emphasis. DMSO-normalized TR-FRET ratios measured for varied doses of Indisulam (black line) and 7 (red line) in DCAF15-DDB1- DDA1-RBM39 recruitment assay. Error bars represent s.d. of the mean from four biological replicates (n=4) in a single experiment. **b**, Effect of 72 h compound 7 (red line) or Indisulam (black line) treatment on viability (CellTiterGlo) of HCT116 cells. Error bars represent s.d. of the mean from four biological replicates (n=4) in a single experiment. Each experiment was performed two independent times. **c**, Western blots showing levels of RBM39 in HCT116 cells following 6 h treatment of indicated concentrations of Indisulam (left), compound 7(right), or DMSO. Data shown from one individual, representative experiment from two independent repeats.

**Supplementary Figure 8.**
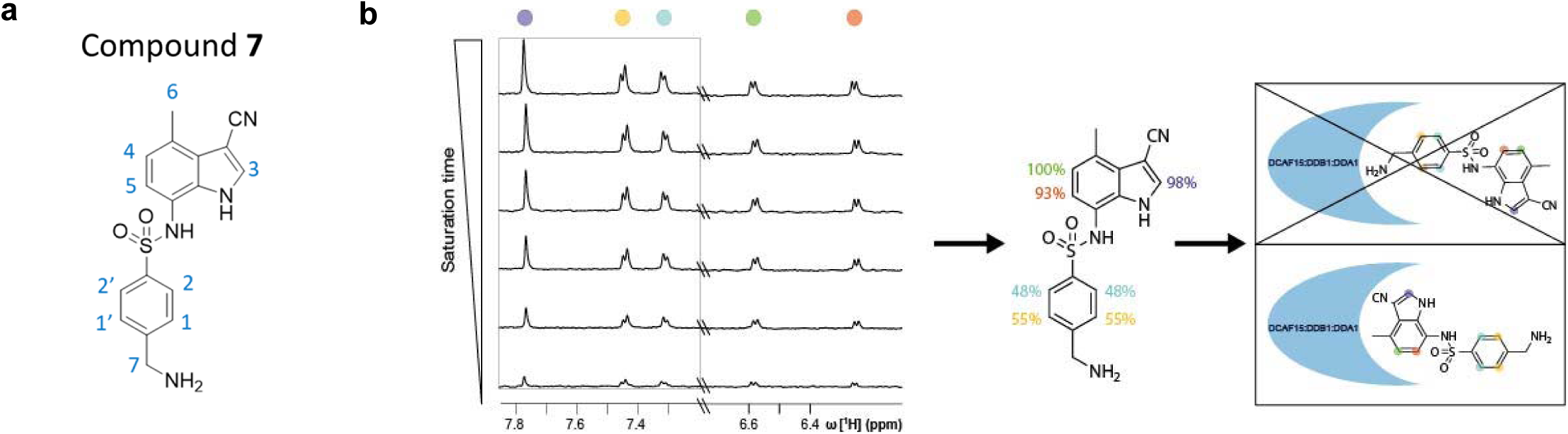
NMR-based epitope mapping of compound 7 highlights regions in proximity to DCAF15. **a**, Structure of the Indisulam analog 7. **b**, ^1^H STD experiments were recorded with different magnetization saturation durations to obtain STD build-up curves (left) which were analyzed to quantify and normalize the amount of magnetization that was transferred from DCAF15-DDB1-DDA1 to each proton in compound 7 (middle). A compound binding model was derived based on the STD results, which show that the protons of the indole receive more magnetization than the ones of the phenyl group and hence, are likely located more closely to the ligase (right).

**Supplementary Figure 9.**
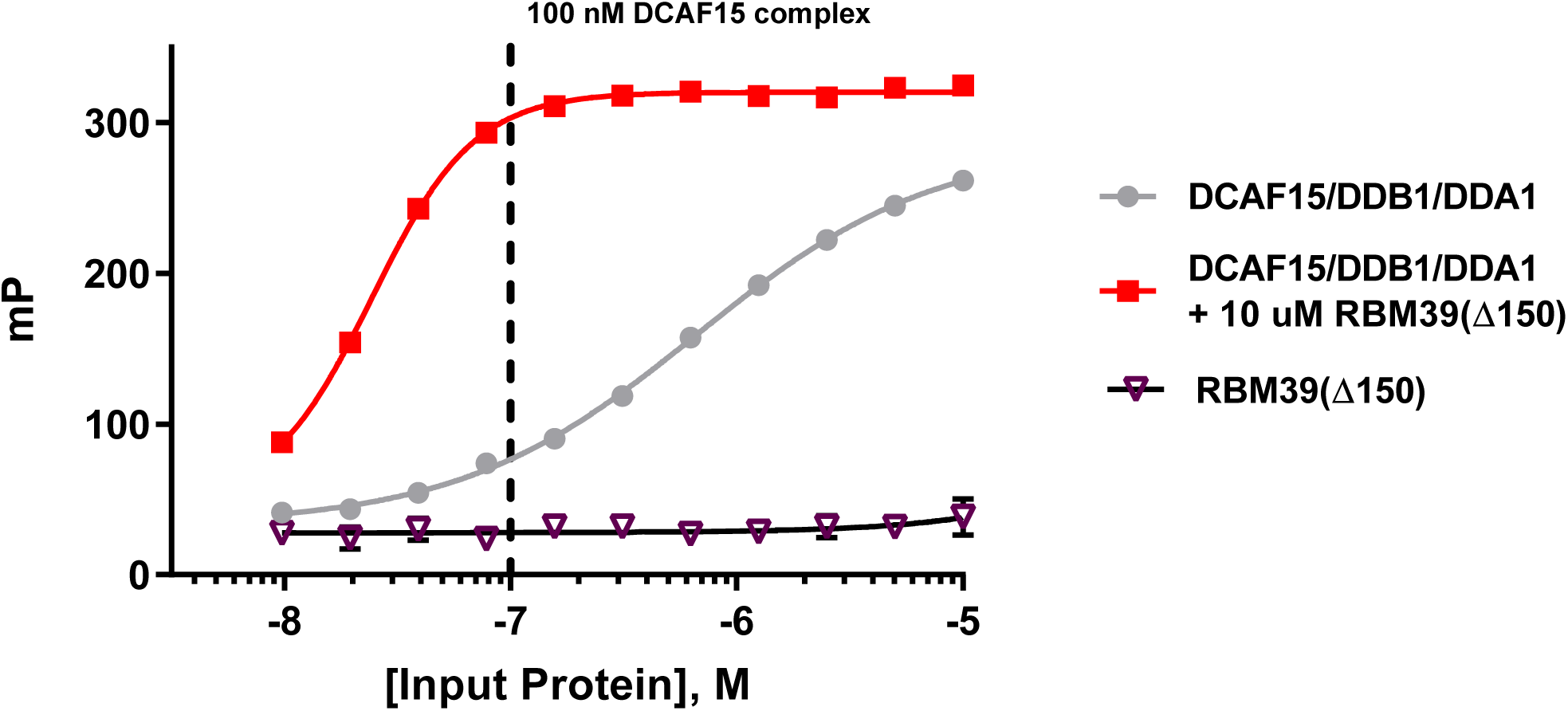
DCAF15-DDB1-DDA1 and RBM39 protein titrations in fluorescence polarization assay with compound 9. Fluorescence polarization (FP) measured for 20 nM compound 9, a FITC-labeled Indisulam analog, following titration of DCAF15-DDB1-DDA1 (grey line) in the presence of 10 µM RBM39(Δ150) (red line), DCAF15-DDB1-DDA1 alone (grey line), or RBM39(Δ150) alone (purple line). While 9 binds DCAF15-DDB1-DDA1 alone, the addition of 10 µM RBM39(Δ150) greatly enhances apparent affinity as 9 forms a ternary complex with both proteins. No binding is detected between 9 and RBM39(Δ150) alone. From these data, low 9 binding occurs at 100 nM DCAF15-DDB1-DDA1, but near maximum binding in the presence of RBM39(Δ150) at the same concentration (vertical dashed line). As this change in signal reflects enhance binding through ternary complex formation between DCAF15-DDB1-DDA1 and RBM39(Δ150), These conditions were chosen to assay impact of RBM39(Δ150) mutation on this phenomenon. Error bars represent s.d. of the mean from four biological replicates (n=8) in a single experiment.

**Supplementary Figure 10.**
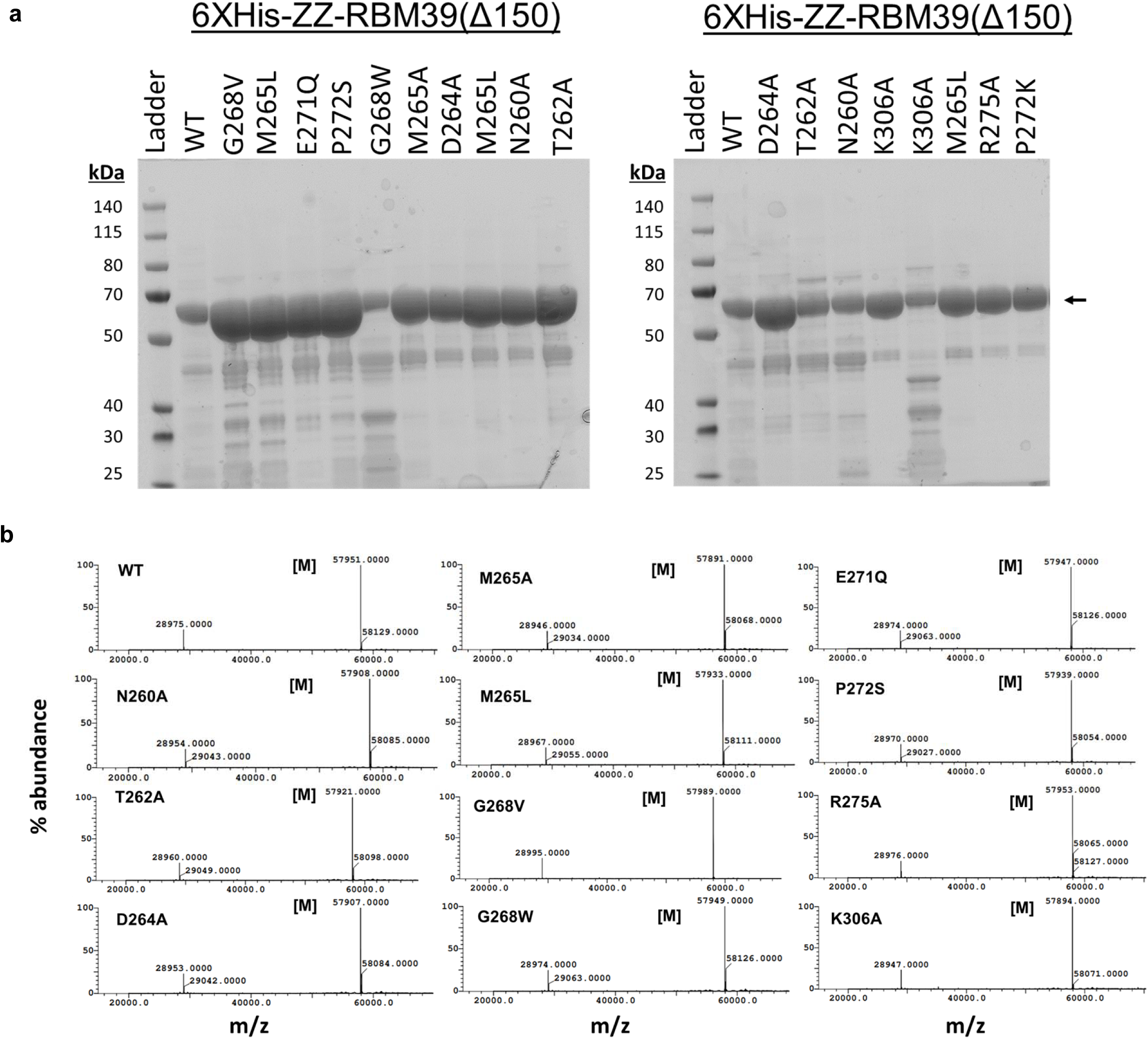
Characterization of purified RBM39 variants. **a**, Coomassie-stained gels of all RBM39(Δ150) variants used in this study and **b**, associated LC-MS analysis and mass determination for each prep. Expected mass is represented by [M].

**Supplementary Figure 11.**
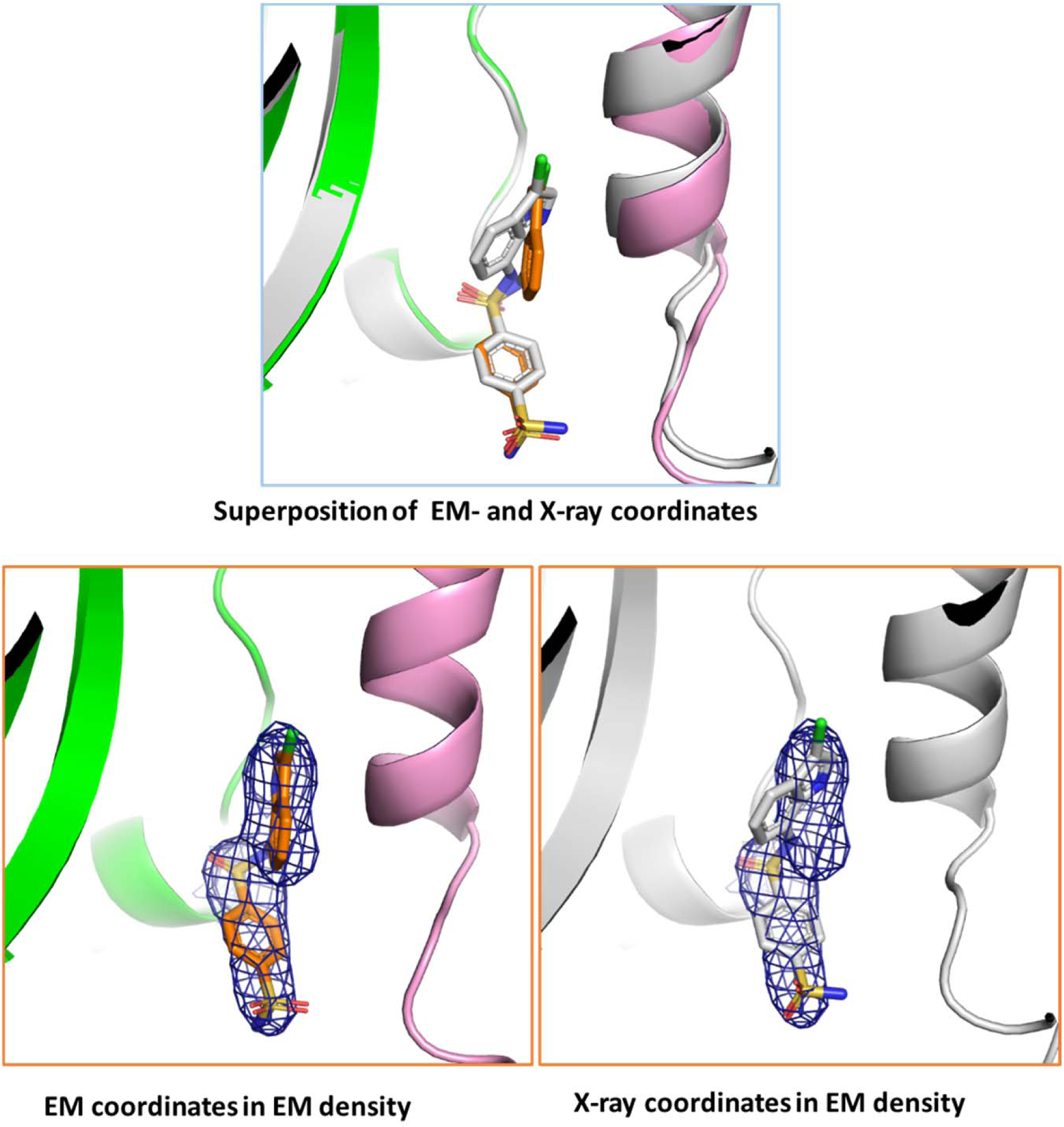
Positioning of Indisulam in Cryo-EM (white) and X-ray (orange) structures. The majority of the Indisulam molecule shows similar conformation (superposition on top), however, the positioning of the chloro-indole group differs by ∼30 degrees between Cryo-EM (bottom left) and X-ray (bottom right) structures. The estimated energy of the indisulam binding pose in the EM structure is ∼1.4 kcal/mol higher versus the X-ray crystal structure assuming the central sulfonamide is deprotonated. However, due to resolution differences in the data used to fit the compounds, it is likely that the structures are approximately equivalent.

## Supplemental discussion

### Differences between the structures of the DCAF15-DDB1-DDA1-RBM39(RRM2)-Indisulam complex by X-ray crystallography and cryo-electron microscopy

The orthogonal determination of the co-structure by two separate structural methods allows both the validation of the determined structures as well as an examination of important differences^1,2^. Generally, the high level of sampling of the macromolecule or macromolecular complex provided by Bragg diffraction coupled with the restriction of movement provided by a three-dimensional crystal matrix allows crystallography to provide higher-resolution data, while the imaging of the macromolecular complex by cryo-EM frozen in vitreous ice allows observation of pertinent dynamics at the cost of some resolution. With the exception of the DDB1-BPB domain, which has been deleted from the crystallographic construct to enforce crystal packing and which cannot be fit in the EM structure due to weak density, the two structures overlap with an RMSD of 1.16 Å between the main-chain atoms of all proteins in the complex, illustrating the high degree of similarity between the two structures (Fig. 2c). However, significant differences do exist. Three loops in DCAF15 which are visible in the crystal structure are not present in the EM structure, and β12 is also not visible. The trajectory of residues 191-214 differs significantly. The N-terminus and C-terminus of RBM39 diverge between the two structures by 4.45 and 8.18 Å, respectively, while the main body of RBM39 is similar. Several loops present in DDB1 in the crystal structure are not visible in the EM structure. Residues 5-69 of DDA1 are visible in the EM structure, while residues 4-76 are visible in the crystal structure. While most of the rotamers of side-chains are similar between the two structures, there are many cases where the side-chains favor different rotamers. Given the differences in resolution of the two structures, it is difficult to explain these results, as the differences in resolution could affect interpretation of the maps and subsequent refinement of the structure. Some of these differences are likely due to the differences in mobility between the two structures. Some could also be affected by the difference in buffer components and pH between the two systems. Additionally, the crystal structure is necessarily determined in the presence of a chemically complex precipitant solution. In general, as strain is not readily accommodated in a crystal matrix, the crystal structure can be viewed as a low-energy state of the quaternary complex, but perhaps not the lowest energy state. The cryo-EM structure can be viewed as a representation of the mobility of the complex free in solution, subject, of course, to the freezing process used in cryo-EM.

The Indisulam binding pocket and the binding pose of Indisulam differs slightly between the two structures (Supplementary Fig. 11). While the majority of the compound shows the same conformation, the position of the chloro-indole group differs by approximately 30 degrees within the hydrophobic pocket. Examination of the pocket reveals that the pocket is slightly larger due to a 1.7 Å shift of both Met265 of RBM39 and Met560 of DCAF15 away from the compound, as well as shift in the rotamer of Val556 from DCAF15. This allows the compound to adopt a binding pose with a slightly lower free energy. Calculations (using Gaussian 2009) estimate the energy of the indisulam binding pose in the co-crystal structure to be ∼1.4 kcal/mol lower in than the binding pose in the EM structure, assuming that the nitrogen in the central phenylsulfonamide is ionized.

## Supplementary Note 1: 2Fo-Fc electron density map for the Indisulam binding pocket and adjacent structure

**Supplementary Note 1:**
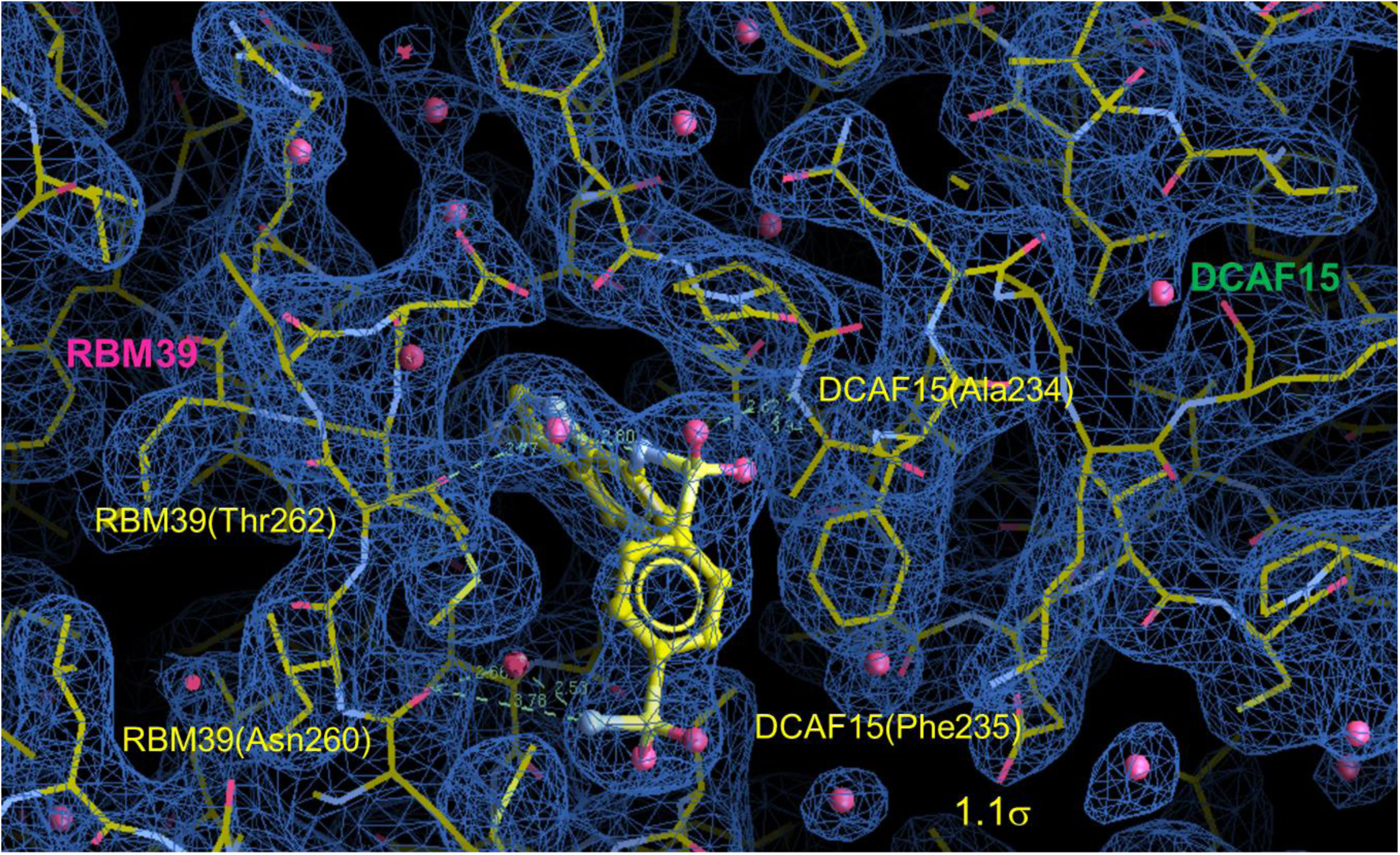
2Fo-Fc electron density map for the Indisulam binding pocket and adjacent structure. The map is shown in blue and is contoured at 1.1 σ. Structural waters are shown and key interactions are shown as dotted lines.

## Supplementary Note 2: Synthesis of Indisulam and analogs

**Indisulam synthesis and characterization**

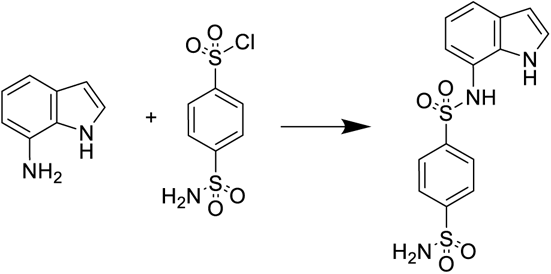

4-sulfamoylbenzene-1-sulfonyl chloride (367 mg, 1.435 mmol) was added to a room temperature solution of 7-aminoindole (187 mg, 1.415 mmol) and pyridine (0.250 mL, 3.09 mmol) in EtOAc (7 mL). The reaction mixture was the stirred at room temperature for 2 hrs. The reaction mixture was diluted with EtOAc (30 mL) then washed with 0.5 M HCl (10 mL), sat. aq. NaHCO3 solution (10 mL), then brine (10 mL). the organic layer was then dried over Na2SO4, filtered, loaded onto Celite and purified over SiO2 column with 0-15% MeOH/DCM to afford N-(1H-indol-7-yl)benzene-1,4-disulfonamide (360mg, 72%). LC-MS: m/z = 352.0297 (M+H^+^); 1H NMR (400 MHz, DMSO-d6) δ 10.79 (s, 1H), 10.12 (s, 1H), 7.93 (s, 4H), 7.55 (s, 2H), 7.36 - 7.29 (m, 2H), 6.83 (t, J = 7.7 Hz, 1H), 6.72 - 6.66 (m, 1H), 6.41 (dd, J = 3.0, 2.0 Hz, 1H).

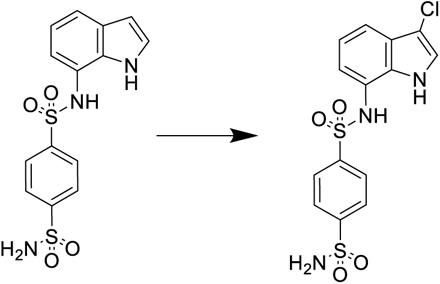

A solution of N-(1H-indol-7-yl)benzene-1,4-disulfonamide (131 mg, 0.373 mmol) in THF (4 mL) was cooled in an ice bath for 10 min. NCS (58mg, 0,42mmol) was then added and stirred in the cold bath for 5 min the removed and stirred at room temperature for 1.5 hr. *ca*. 10% conversion is seen. 1 drop of conc. HCl was then added and stirred at room temperature for 30 min. The reaction mixture was diluted with water (10 mL) and EtOAc (15 mL). The mixture was separated and the aqueous layer was extracted with EtOAc (10 mL). The combined organic layers were then washed with sat. aq. NaHCO3 solution (10 mL) then brine (10 mL). The organic layer was then dried over Na2SO4, filtered, loaded onto celite and purified over SiO2 with 0-100% EtOAc/heptane to afford Indisulam (93mg, 63%). LC-MS: m/z = 386.0037 (M+H^+^); ^1^H NMR (400 MHz, DMSO-d6) δ 11.10 (s, 1H), 10.21 (s, 1H), 7.97 - 7.90 (m, 4H), 7.56 (s, 2H), 7.50 (d, J = 2.7 Hz, 1H), 7.29 (d, J = 7.9 Hz, 1H), 6.96 (t, J = 7.8 Hz, 1H), 6.76 (d, J = 7.1 Hz, 1H).

**General scheme for synthesis of compounds 1, 2, and 4**

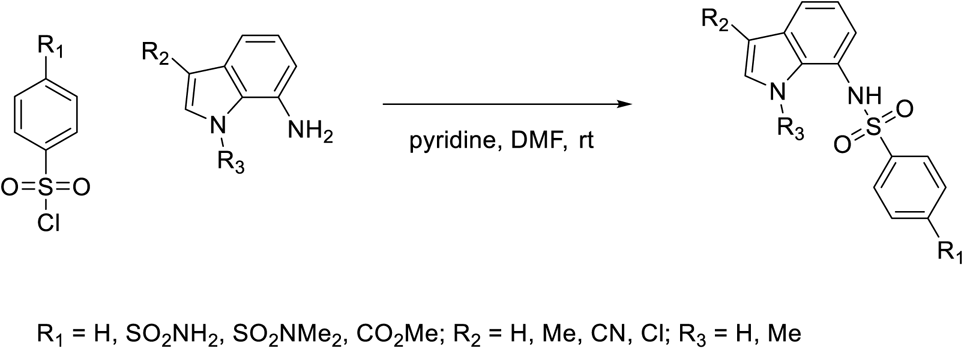

**Synthesis and characterization of compounds 1, 2, and 4**

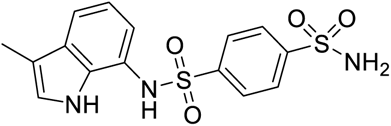

**Compound 1** (N-(3-methyl-1H-indol-7-yl)benzene-1,4-disulfonamide)

3-methyl-1H-indol-7-amine (15 mg, 0.103 mmol) was suspended in Pyridine (Volume: 1 mL) and 4-sulfamoylbenzenesulfonyl chloride (34.1 mg, 0.133 mmol) was added. The reaction mixture was stirred at room temp for 16 hrs. Pyridine was removed by vacuo and redissolved with DMSO. Subjected to prep HPLC and the fractions were lyophilyzed to yield the desired compound.

^1^H NMR (400 MHz, DMSO-*d6*) δ 10.34 (s, 1H), 10.01 (s, 1H), 7.86 (brs, 4H), 7.47 (s, 2H), 7.20 (dt, *J* = 7.8, 0.8 Hz, 1H), 7.02 (dd, *J* = 2.5, 1.2 Hz, 1H), 6.76 (t, *J* = 7.7 Hz, 1H), 6.66 (dd, *J* = 7.6, 1.0 Hz, 1H), 2.13 (s, 3H). LC-MS: m/z 366.4 [M+1].

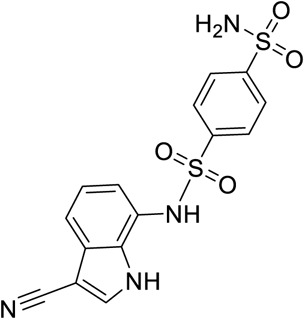

**Compound 2** (N-(3-cyano-1H-indol-7-yl)benzene-1,4-disulfonamide)

A mixture of 7-amino-1H-indole-3-carbonitrile (16 mg, 0.102 mmol) and sulphonyl chloride (-, 0.112 mmol) in DMF (Volume: 1 ml, Ratio: 5.00, Total Volume: 6.00 ml) and pyridine (Volume: 0.2 ml, Ratio: 1.000, Total Volume: 1.200 ml) was stirred at RT overnight. The reaction was purified reverse phase HPLC to give the product as off white solids.

^1^H NMR (400 MHz, Methanol-*d4*) δ 7.99 - 7.96 (m, 2H), 7.96 (s, 1H), 7.83 - 7.78 (m, 2H), 7.52 (dd, J = 8.0, 0.9 Hz, 1H), 7.04 (t, J = 7.8 Hz, 1H), 6.60 (dd, J = 7.6, 0.9 Hz, 1H). LC-MS: m/z 377.1 [M+1].

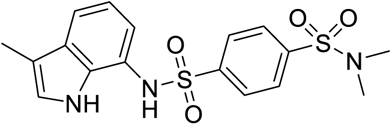

**Compound 4** (*N^1^*,*N*^1^-dimethyl-*N*^4^-(3-methyl-1H-indol-7-yl)benzene-1,4-disulfonamide)

3-methyl-1H-indol-7-amine (10 mg, 0.068 mmol) was dissolved in Pyridine (Volume: 1 mL) and 4-(N,N-dimethylsulfamoyl)benzenesulfonyl chloride (23.29 mg, 0.082 mmol) was added. The mixture was stirred for 16hrs. Pyridine was removed and dried. Purified by 3 silica gel chromatogaphyby using DCM/MeOH to yield final product (13 mg, 0.031 mmol, 45.9 % yield).

^1^H NMR (400 MHz, DMSO-*d_6_*) δ 10.25 (s, 1H), 9.97 (s, 1H), 7.87-7.80 (m, 2H), 7.79-7.73 (m, 2H), 7.21 (dt, *J* = 7.9, 0.8 Hz, 1H), 6.97 (dd, *J* = 2.5, 1.2 Hz, 1H), 6.77 (t, *J* = 7.7 Hz, 1H), 6.61 (dd, *J* = 7.6, 1.0 Hz, 1H), 2.49 (s, 6H), 2.11 (s, 3H). LC-MS: m/z 394.2 [M+1].

**Compound 3 synthesis and characterization**

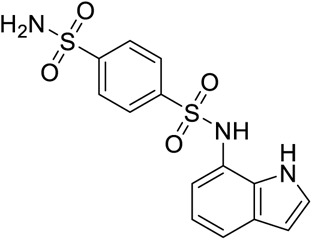

**Compound 3** (N-(1H-indol-7-yl)benzene-1,4-disulfonamide)^14^

4-sulfamoylbenzene-1-sulfonyl chloride (367 mg, 1.4 mmol) was added to a room temperature solution of 7-aminoindole (187 mg, 1.4 mmol) and pyridine (0.250 mL, 3.09 mmol) in EtOAc (7 mL). Te reaction mixture was then stirred at room temperature for 2 hrs. The reaction mixture was diluted with EtOAc (30 mL) then washed with 0.5 M HCl (10 mL), sat. aq. NaHCO3 solution (10 mL), then brine (10 mL). The organic layer was then dried over Na2SO4, filtered, and concentrated to dryness affording a brown amorphous solid. The solid was then purified by silica gel chromatography, eluting with 0-15% MeOH/DCM, to yield an orange solid. The solid was then triturated with Et2O/heptane. The mixture was filtered then washed several times with heptane. The solid was then dried under vacuum filtration to afford the desired product as a light pink solid (360 mg, 1.01 mmol, 72% yield. 1H NMR (400 MHz, DMSO-d6) δ 10.79 (s, 1H), 10.12 (s, 1H), 7.93 (s, 4H), 7.55 (s, 2H), 7.36 - 7.29 (m, 2H), 6.83 (t, J = 7.7 Hz, 1H), 6.72 - 6.66 (m, 1H), 6.41 (dd, J = 3.0, 2.0 Hz, 1H). LC-MS: m/z 351.9 [M+H]. HRMS (M+H) calculated C14H14N3O4S2 352.0426, found 352.0297.

**Compound 5 synthesis and characterization**

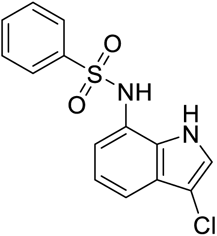

**Compound 5** (N-(3-chloro-1H-indol-7-yl)benzenesulfonamide)

Et3N (1.8 mmol) was added to a *ca*. 0 °C suspension of 3-chloro-1H-indol-7-amine ^15^ in DCM (10 mL). Benzenesulfonyl chloride (160 mg, 0.9 mmol) was then added and the reaction mixture was allowed to stir and warm up to room temperature overnight. The reaction mixture was quenched with ice water then extracted with DCM (2 X 10 mL). The combined organic layers were washed with water then brine. The organic layer was then dried over Na2SO4, filtered, and concentrated to dryness. The crude material was then purified by silica gel chromotography, eluting with 20-25% EtOAc/hexane, to afford the desired product as a brown solid (28 mg, 0.09 mmol, 10% yield).

1H NMR (300 MHz, DMSO-d6): δ 11.01 (s, 1H), 9.99 (s, 1H), 7.74-7.71 (m, 2H), 7.62-7.57 (m, 1H), 7.53-7.46 (m, 3H), 7.24 (d, J = 9 Hz, 1H), 6.94 (t, J = 10 Hz, 1H), 6.77 (d, J = 9.6 Hz, 1H). LC-MS: m/z 305.2 [M-H]

**Compound 8 synthesis and characterization**

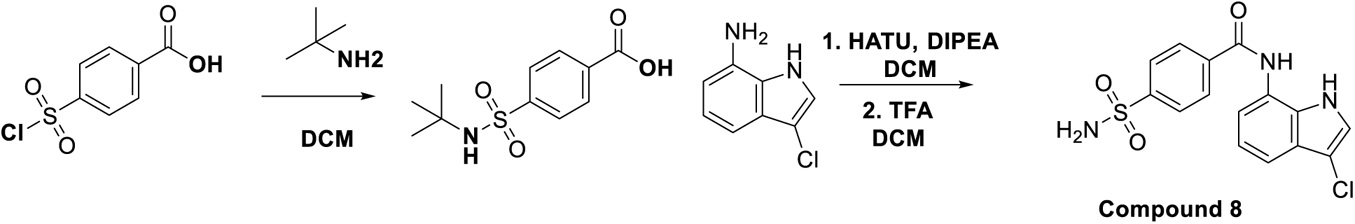

**Compound 8** (N-(3-chloro-1H-indol-7-yl)-4-sulfamoylbenzamide)

Step 1: To a stirred solution of 4-(chlorosulfonyl)benzoic acid (3g, 13.6 mmol) in DCM (30 mL) was added a solution of tert-butyl amine (5.7 mL, 54 mmol) in DCM (20 mL) at 0 °C. The reaction mixture was allowed to stir and warm up to room temperature over 3 hrs. The reaction mixture was filtered and the solid was washed wtih DCM then dried under vacuum filtration. The solid was collected then put into water then slowly and carefully acidified to *ca*. pH 3-4 using 5N HCl. The resulting suspension was stirred at room temperature for 20 min then filtered. The solid was washed with water then dried under vacuum filtration to afford the desired 4-(N-(tert-butyl)sulfamoyl)benzoic acid as a white solid (2.5 g, 9.7 mmol, 71% yield). LC-MS: m/z = 255.9 [M-H].

Step 2: To a solution of 3-chloro-1H-indol-7-amine^15^ (200 mg, 1.2 mmol) in DMF (7 mL) were added 4-(N-(tert-butyl)sulfamoyl)benzoic acid (308 mg, 1.2 mmol), HATU (684 mg, 1.8 mmol), and DIPEA (1.07 mL, 6 mmol). The reaction mixture was then stirred at room temperature for 16 hrs. The reaction mixture was concentrated under reduced pressure to remove DMF and the crude material was purified *via* silica gel chromotagraphy, eluting with 30% EtOAc/heptane, to afford the tert-butyl protected compound 8 as a brown solid (480 mg, 98%). LC-MS: m/z 406.0 [M+H].

Step 3: To a stirred solution of the brown solid (400 mg, 0.27 mmol) from step 2 in DCM (5 mL) was added TFA (2 mL) at 0 °C. The reaction was then stirred at room temperature for 16 hrs. The reaction mixture was quenched with sat. aq. NaHCO3 solution then diluted with water. The mixture was extracted with DCM (2 X 10 mL) and the combined organic layers were concentrated to dryness *in vacuo*. The resulting residue was purified by silica gel chromatogaphy, eluting with 70% EtOAc/hexane, to afford the desired COMPOUND 8 as brown solid (20 mg, 0.05 mmol, 19% yield).

1H NMR (300 MHz, DMSO-d6): δ 11.2 (br s, 1H), 10.4 (s, 1H), 8.19 (d, J = 11.6 Hz, 2H), 7.98 (d, J = 11.6 Hz, 2H), 7.56-7.55 (m, 3H), 7.40 (t, J = 10 Hz, 2H), 7.14 (t, J = 10 Hz, 1H). MS: m/z 347.9 [M+H]

**Compound 7 synthesis and characterization**

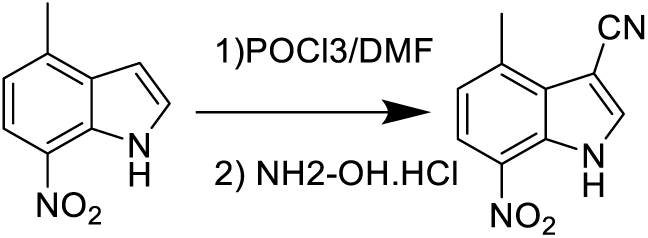

To 1 mL of dimethylformamide was added POCl3 (0.317 mL, 3.41 mmol) at 0° C., followed by stirring at 0° C. for 0.5 hour. To the reaction mixture was then added a solution of 4- methyl-7-nitro-1H-indole (500 mg, 2.84 mmol) in 2.0ml DMF at 0° C., followed by heating and stirring at 60° C. for 2 hours. To the reaction mixture was then added dropwise a solution of hydroxylamine hydrochloride (394 mg, 5.68 mmol) in 3.0ml DMF with keeping the internal temperature below 80° C., followed by heating and stirring at 60° C. for 40 minutes. The reaction was cooled in an ice bath and 18ml ice water was added to the reaction mixture, which was further stirred 1hr. The precipitated crystals were collected by filtration and washed with water. The crystals were suspended in 18ml H_2_O, 1N NaOH was added to the suspension to adjust pH to 7, and then the crystals were collected by filtration, washed with water and dried to afford 4-methyl-7-nitro-1H-indole-3-carbonitrile (520mg, 91%). The crude product was used as it was in the next step. LC-MS m/z = 200.2 (M-H^+^).

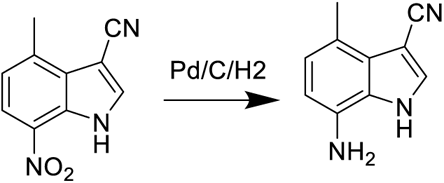

To a solution of 4-methyl-7-nitro-1H-indole-3-carbonitrile (520 mg, 2.58 mmol) in MeOH (Volume: 10 mL, Ratio: 1.000) and THF (Volume: 10 mL, Ratio: 1.000), Pd/C (138 mg, 0.129 mmol) was added. The reaction mixture was treated under H2 ballon at rt for 2hr. The reaction was filtered and washed with acetone. The filtrated was concentrated to afford 7-amino-4-methyl-1H-indole-3-carbonitrile (443mg, 100%). The crude solid was used as it was in the next step. LC-MS: m/z = 170.0 (M-H^-^).

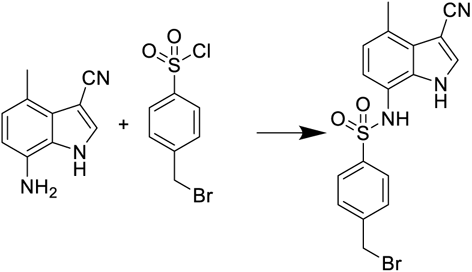

To a solution of 7-amino-4-methyl-1H-indole-3-carbonitrile (300 mg, 1.752 mmol) in THF (Volume: 12 mL) at 0oC, pyridine (0.354 mL, 4.38 mmol) was added. After 10min. at 0oC, 4-bromomethyl benzenesulphonyl chloride (661 mg, 2.453 mmol) was added. The reaction stirred at rt overnight. The reaction was extracted between H_2_O and ethylacetate. Combined all the organics, dried, concentrated and purified over SiO2 with 40% ethylacetate/heptane to afford 4-(bromomethyl)-N-(3-cyano-4-methyl-1H-indol-7- yl)benzenesulfonamide (250mg, 35%). LC-MS: m/z = 402.0 (M-H^-^).

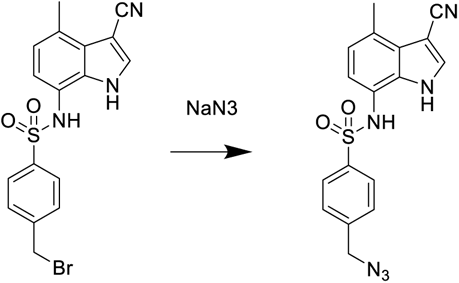

To solution of 4-(bromomethyl)-N-(3-cyano-4-methyl-1H-indol-7-yl)benzenesulfonamide (250 mg, 0.618 mmol) in DMF (Volume: 2 mL), NaN3 (201 mg, 3.09 mmol) was added, followed by TBAI (45.7 mg, 0.124 mmol). The reaction was treated at 120oC for 0.5hr under microwave. The reaction was extracted between H_2_O and ethylacetate. Combined all the organics, dried, concentrated and purified over SiO2 with 40% ethylacetate/heptane to afford 4-(azidomethyl)-N-(3-cyano-4-methyl-1H-indol-7-yl)benzenesulfonamide (200mg, 88%). LC-MS: m/z = 365.1 (M-H^+^).

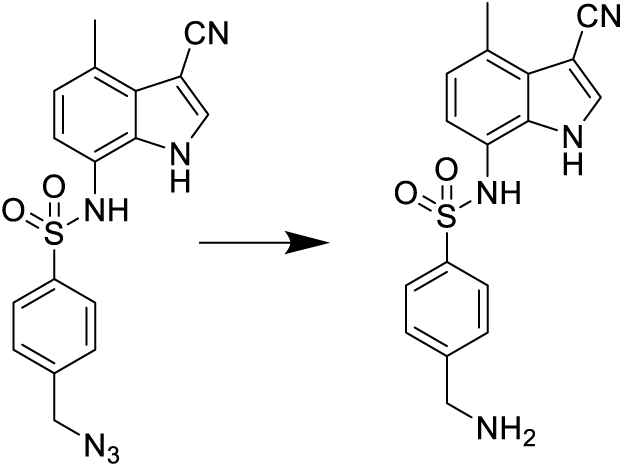

To the solution of 4-(azidomethyl)-N-(3-cyano-4-methyl-1H-indol-7- yl)benzenesulfonamide (680 mg, 1.856 mmol) in THF (Volume: 10 mL, Ratio: 10.00) and H_2_O (Volume: 1 mL, Ratio: 1.000), triphenylphosphine (730 mg, 2.78 mmol) was added. The reaction was treated at 60oC for 3hr. The reaction was extracted between H_2_O and ethylacetate. All organics were combined, dried, concentrated and purified over SiO2 with 10% MeOH/DCM to afford 4-(aminomethyl)-N-(3-cyano-4-methyl-1H-indol-7- yl)benzenesulfonamide (500 mg, 79%). LC-MS: m/z = 341.2 (M+H^+^).

**Compound 9 synthesis and characterization**

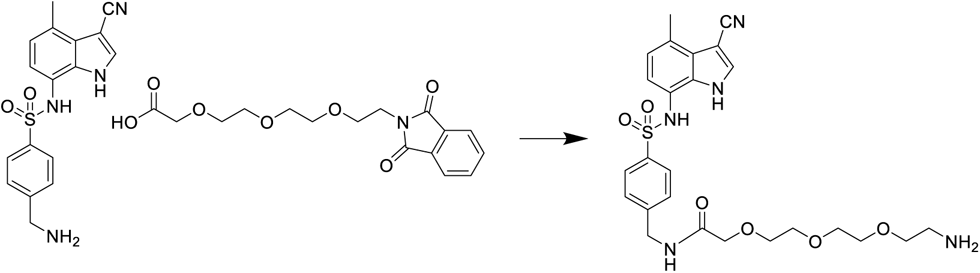

To a solution of 4-(aminomethyl)-N-(3-cyano-4-methyl-1H-indol-7- yl)benzenesulfonamide (40 mg, 0.118 mmol) in DMF (Volume: 1.0 mL) at 0 ⁰C, 2-(2-(2- (2-(1,3-dioxoisoindolin-2-yl)ethoxy)ethoxy)ethoxy)acetic acid (59.5 mg, 0.176 mmol) and DIPEA (0.051 mL, 0.294 mmol) were added, followed by HATU (89 mg, 0.235 mmol). The reaction was stirred at 0 ^⁰^C for 15min. Next, hydrazine (0.037 mL, 1.175 mmol) was added and the reaction was stirred at 50 ⁰C for 0.5hr. The reaction was diluted with DMSO and purified over RP HPLC under basic condition with the detection of the desired product MW to afford 2-(2-(2-(2-aminoethoxy)ethoxy)ethoxy)-N-(4-(N-(3-cyano-4-methyl-1H- indol-7-yl)sulfamoyl)benzyl)acetamide (25mg, 40%). LC-MS: m/z = 530.18 (M+H^+^).

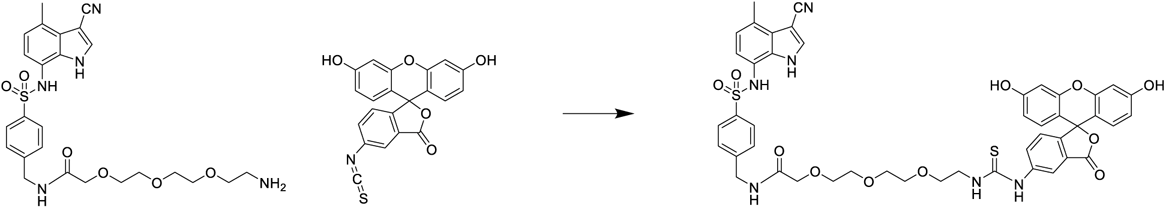

To a solution of 2-(2-(2-(2-aminoethoxy)ethoxy)ethoxy)-N-(4-(N-(3-cyano-4-methyl-1H- indol-7-yl)sulfamoyl)benzyl)acetamide (11 mg, 0.021 mmol) in DMF (Volume: 1.0 mL), FITC (9.70 mg, 0.025 mmol) and DIPEA (10.88 µl, 0.062 mmol) were added. The reaction was stirred at rt for overnight. The reaction was diluted with DMSO and purified over RP HPLC under acidic condition with UV detection to afford the desired product (15mg, 75%). LC-MS: 919.5 (M+H^+^).

## References

1. Bondeson, D. P. & Crews, C. M. Targeted Protein Degradation by Small Molecules. Annu. Rev. Pharmacol. Toxicol. (2017). doi:10.1146/annurev-pharmtox-010715-103507

2. Collins, I., Wang, H., Caldwell, J. J. & Chopra, R. Chemical approaches to targeted protein degradation through modulation of the ubiquitin–proteasome pathway. Biochem. J. (2017). doi:10.1042/bcj20160762

3. Ciechanover, A. Intracellular protein degradation: From a Vague Idea, through the lysosome and the ubiquitin-proteasome system, and onto human diseases and drug targeting (Nobel Lecture). Angewandte Chemie - International Edition 44, 5944–5967 (2005).

4. Thrower, J. S. Recognition of the polyubiquitin proteolytic signal. EMBO J. (2000). doi:10.1093/emboj/19.1.94

5. Yu, H. & Matouschek, A. Recognition of Client Proteins by the Proteasome. Annu. Rev. Biophys. (2017). doi:10.1146/annurev-biophys-070816-033719

6. Finley, D., Chen, X. & Walters, K. J. Gates, Channels, and Switches: Elements of the Proteasome Machine. Trends in Biochemical Sciences (2016). doi:10.1016/j.tibs.2015.10.009

7. Neutzner, M. & Neutzner, A. Enzymes of ubiquitination and deubiquitination. Essays Biochem. (2012). doi:10.1042/bse0520037

8. Zheng, N. & Shabek, N. Ubiquitin Ligases: Structure, Function, and Regulation. Annu. Rev. Biochem. (2017). doi:10.1146/annurev-biochem-060815-014922

9. Wu, Y. L. et al. Structural basis for an unexpected mode of SERM-Mediated ER antagonism. Mol. Cell (2005). doi:10.1016/j.molcel.2005.04.014

10. Patel, H. K. & Bihani, T. Selective estrogen receptor modulators (SERMs) and selective estrogen receptor degraders (SERDs) in cancer treatment. Pharmacology and Therapeutics (2018). doi:10.1016/j.pharmthera.2017.12.012

11. Neklesa, T. K., Winkler, J. D. & Crews, C. M. Targeted protein degradation by PROTACs. Pharmacology and Therapeutics (2017). doi:10.1016/j.pharmthera.2017.02.027

12. Larrieu, A. & Vernoux, T. Comparison of plant hormone signalling systems. Essays Biochem. (2015). doi:10.1042/bse0580165

13. Tan, X. et al. Mechanism of auxin perception by the TIR1 ubiquitin ligase. Nature (2007). doi:10.1038/nature05731

14. Chamberlain, P. P. & Cathers, B. E. Cereblon modulators: Low molecular weight inducers of protein degradation. Drug Discovery Today: Technologies (2019). doi:10.1016/j.ddtec.2019.02.004

15. Supuran, C. T. Indisulam: an anticancer sulfonamide in clinical development. Expert Opin. Investig. Drugs (2003). doi:10.1517/eoid.12.2.283.21409

16. Ozawa, Y. et al. E7070, a novel sulphonamide agent with potent antitumour activity in vitro and in vivo. Eur. J. Cancer (2001). doi:10.1016/S0959-8049(01)00275-1

17. Uehara, T. et al. Selective degradation of splicing factor CAPERα by anticancer sulfonamides. Nat. Chem. Biol. (2017). doi:10.1038/nchembio.2363

18. Han, T. et al. Anticancer sulfonamides target splicing by inducing RBM39 degradation via recruitment to DCAF15. Science *(80-.).* (2017). doi:10.1126/science.aal3755

19. Laskowski, R. A., Jabłońska, J., Pravda, L., Vařeková, R. S. & Thornton, J. M. PDBsum: Structural summaries of PDB entries. Protein Sci. 27, 129–134 (2018).

20. Holm, L. & Rosenstrï¿½m, P. Dali server: conservation mapping in 3D. Nucleic Acids Res. 38, W545–W549 (2010).

21. Dias, J. et al. Structural analysis of the KANSL1/WDR5/ KANSL2 complex reveals that WDR5 is required for efficient assembly and chromatin targeting of the NSL complex. Genes Dev. (2014). doi:10.1101/gad.240200.114

22. Wysocka, J. et al. WDR5 associates with histone H3 methylated at K4 and is essential for H3 K4 methylation and vertebrate development. Cell (2005). doi:10.1016/j.cell.2005.03.036

23. Song, J. J. & Kingston, R. E. WDR5 interacts with mixed lineage leukemia (MLL) protein via the histone H3-binding pocket. J. Biol. Chem. (2008). doi:10.1074/jbc.M806900200

24. Qu, Q. et al. Structure and Conformational Dynamics of a COMPASS Histone H3K4 Methyltransferase Complex. Cell 174, 1117–1126.e12 (2018).

25. Jain, B. P. & Pandey, S. WD40 Repeat Proteins: Signalling Scaffold with Diverse Functions. Protein J. 37, 391–406 (2018).

26. Xue, B., Dunbrack, R. L., Williams, R. W., Dunker, A. K. & Uversky, V. N. PONDR-FIT: a meta-predictor of intrinsically disordered amino acids. Biochim. Biophys. Acta 1804, 996–1010 (2010).

27. Wu, Y. et al. The DDB1–DCAF1–Vpr–UNG2 crystal structure reveals how HIV-1 Vpr steers human UNG2 toward destruction. Nat. Struct. Mol. Biol. 23, 933– 940 (2016).

28. Scrima, A. et al. Structural Basis of UV DNA-Damage Recognition by the DDB1-DDB2 Complex. Cell (2008). doi:10.1016/j.cell.2008.10.045

29. Fischer, E. S. et al. Structure of the DDB1-CRBN E3 ubiquitin ligase in complex with thalidomide. Nature (2014). doi:10.1038/nature13527

30. Shabek, N. et al. Structural insights into DDA1 function as a core component of the CRL4-DDB1 ubiquitin ligase. Cell Discov 4, 67–67 (2018).

31. Chambers, J. C., Kenan, D., Martin, B. J. & Keene, J. D. Genomic structure and amino acid sequence domains of the human La autoantigen. J. Biol. Chem. 263, 18043–51 (1988).

32. Dreyfuss, G., Swanson, M. S. & Piñol-Roma, S. Heterogeneous nuclear ribonucleoprotein particles and the pathway of mRNA formation. Trends Biochem. Sci. 13, 86–91 (1988).

33. Murray, J. M. & Bussiere, D. E. Targeting the Purinome. in *Methods in molecular biology (Clifton*, N.J.) 575, 47–92 (2009).

34. Molecular Operating Environment (MOE), 2013.08. Molecular Operating Environment (MOE), 2013.08; Chemical Computing Group Inc., 1010 Sherbooke St. West, Suite #910, Montreal, QC, Canada, H3A 2R7. Mol. Oper. Environ. (MOE), 2013.08; Chem. Comput. Gr. Inc., *1010 Sherbooke St. West, Suite #910, Montr. QC, Canada, H3A 2R7*, *2013.* (2016).

35. Milletti, F., Storchi, L., Sforna, G. & Cruciani, G. New and original pKa prediction method using grid molecular interaction fields. J. Chem. Inf. Model. (2007). doi:10.1021/ci700018y

36. Chardin, P. & McCormick, F. Brefeldin A: The advantage of being uncompetitive. Cell (1999). doi:10.1016/S0092-8674(00)80724-2

37. Huh, K. et al. Human Papillomavirus Type 16 E7 Oncoprotein Associates with the Cullin 2 Ubiquitin Ligase Complex, Which Contributes to Degradation of the Retinoblastoma Tumor Suppressor. J. Virol. (2007). doi:10.1128/jvi.00881-07

38. Martinez-Zapien, D. et al. Structure of the E6/E6AP/p53 complex required for HPV-mediated degradation of p53. Nature (2016). doi:10.1038/nature16481

39. Poirson, J. et al. Mapping the interactome of HPV E6 and E7 oncoproteins with the ubiquitin-proteasome system. FEBS Journal (2017). doi:10.1111/febs.14193

40. Pick, E. et al. Mammalian DET1 Regulates Cul4A Activity and Forms Stable Complexes with E2 Ubiquitin-Conjugating Enzymes. Mol. Cell. Biol. (2007). doi:10.1128/mcb.02432-06

41. Olma, M. H. et al. An interaction network of the mammalian COP9 signalosome identifies Dda1 as a core subunit of multiple Cul4-based E3 ligases. J. Cell Sci. (2009). doi:10.1242/jcs.043539

42. Gao, S. et al. Activation of c-Abl kinase potentiates the anti-myeloma drug lenalidomide by promoting DDA1 protein recruitment to the CRL4 ubiquitin ligase. J. Biol. Chem. (2017). doi:10.1074/jbc.M116.761551

43. Zhu, Y. X. et al. Identification of cereblon-binding proteins and relationship with response and survival after IMiDs in multiple myeloma. Blood (2014). doi:10.1182/blood-2014-02-557819

44. Krönke, J. et al. Lenalidomide causes selective degradation of IKZF1 and IKZF3 in multiple myeloma cells. Science *(80-.).* (2014). doi:10.1126/science.1244851

45. Sievers, Q. L. et al. Defining the human C2H2 zinc finger degrome targeted by thalidomide analogs through CRBN. Science *(80-.).* (2018). doi:10.1126/science.aat0572

46. Petzold, G., Fischer, E. S. & Thomä, N. H. Structural basis of lenalidomide-induced CK1α degradation by the CRL4 CRBN ubiquitin ligase. Nature (2016). doi:10.1038/nature16979

47. Matyskiela, M. E. et al. A novel cereblon modulator recruits GSPT1 to the CRL4 CRBN ubiquitin ligase. Nature (2016). doi:10.1038/nature18611

48. Laue, T. M., Shah, B. D., Ridgeway, T. M. & Pelletier, S. L. Computer-aided interpretation of analytical sedimentation data for proteins. in Analytical ultracentrifugation in biochemistry and polymer science (1992).

49. Schuck, P. Size-distribution analysis of macromolecules by sedimentation velocity ultracentrifugation and Lamm equation modeling. Biophys. J. (2000). doi:10.1016/S0006-3495(00)76713-0

50. Mayer, M. & Meyer, B. Group epitope mapping by saturation transfer difference NMR to identify segments of a ligand in direct contact with a protein receptor. J. Am. Chem. Soc. (2001). doi:10.1021/ja0100120

51. Bhunia, A., Bhattacharjya, S. & Chatterjee, S. Applications of saturation transfer difference NMR in biological systems. Drug Discovery Today (2012). doi:10.1016/j.drudis.2011.12.016

52. Combe, C. W., Fischer, L. & Rappsilber, J. xiNET: Cross-link Network Maps With Residue Resolution. Mol. Cell. Proteomics (2015). doi:10.1074/mcp.o114.042259

53. Adams, P. D. et al. PHENIX: A comprehensive Python-based system for macromolecular structure solution. Acta Crystallogr. Sect. D Biol. Crystallogr. (2010). doi:10.1107/S0907444909052925

54. Blanc, E. et al. Refinement of severely incomplete structures with maximum likelihood in BUSTER-TNT. Acta Crystallogr. Sect. D Biol. Crystallogr. (2004). doi:10.1107/S0907444904016427

55. Chen, V. B. et al. *MolProbity* : all-atom structure validation for macromolecular crystallography. Acta Crystallogr. Sect. D Biol. Crystallogr. 66, 12–21 (2010).

56. Grant, T., Rohou, A. & Grigorieff, N. CisTEM, user-friendly software for single-particle image processing. Elife (2018). doi:10.7554/eLife.35383

57. Zivanov, J. et al. New tools for automated high-resolution cryo-EM structure determination in RELION-3. Elife (2018). doi:10.7554/eLife.42166

58. Emsley, P., Lohkamp, B., Scott, W. G. & Cowtan, K. Features and development of Coot. Acta Crystallogr. Sect. D Biol. Crystallogr. (2010). doi:10.1107/S0907444910007493

59. Erb, M. A. et al. Transcription control by the ENL YEATS domain in acute leukaemia. Nature (2017). doi:10.1038/nature21688

60. Wessel, D. & Flügge, U. I. A method for the quantitative recovery of protein in dilute solution in the presence of detergents and lipids. Anal. Biochem. (1984). doi:10.1016/0003-2697(84)90782-6

## References

1. Wang, H.-W. & Wang, J.-W. How cryo-electron microscopy and X-ray crystallography complement each other. Protein Sci. 26, 32–39 (2017).

2. Vénien-Bryan, C., Li, Z., Vuillard, L. & Boutin, J. A. Cryo-electron microscopy and X-ray crystallography: complementary approaches to structural biology and drug discovery. Acta Crystallogr. Sect. F Struct. Biol. Commun. 73, 174–183 (2017).

